# Cone Opponent Functional Domains in Primary Visual Cortex Combine Signals for Color Appearance Mechanisms

**DOI:** 10.1101/2020.09.22.309054

**Authors:** Peichao Li, Anupam K. Garg, Li A. Zhang, Mohammad S. Rashid, Edward M. Callaway

## Abstract

Studies of color perception have led to mechanistic models of how cone-opponent signals from retinal ganglion cells are integrated to generate color appearance. But it is unknown how this hypothesized integration occurs in the brain. Here we show that cone-opponent signals transmitted from retina to primary visual cortex (V1) are integrated through highly organized circuits within V1 to implement the color opponent interactions required for color appearance. Combining intrinsic signal optical imaging (ISI) and 2-photon calcium imaging (2PCI) at single cell resolution, we demonstrate cone-opponent functional domains (COFDs) that combine L/M cone-opponent and S/L+M cone-opponent signals following the rules predicted from psychophysical studies of color perception. These give rise to an orderly organization of hue preferences of the neurons within the COFDs and the generation of hue “pinwheels”. Thus, spatially organized neural circuits mediate an orderly transition from cone-opponency to color appearance that begins in V1.

---

Color vision research has a long history of rigorous psychophysical investigation to develop mechanistic models. This has led to some of the greatest successes in linking neural circuit mechanisms to perceptual phenomena. Psychophysical studies led to a Three-stage model^1–5^ (Fig. 1) that presaged the later discovery of three cone types (Stage 1: L-long, M-middle, and S-short wavelength sensitive) and four cone-opponent mechanisms (Stage 2: L+M-, M+L-, S+(L+M)-, and (L+M)+S-). We now know that these cone-opponent mechanisms are generated by precise retinal circuits to give rise to four types of L/M opponent neurons and two types of S/(L+M) opponent neurons^6–8^. These retinal ganglion cell types connect to the primary visual cortex (V1) via the lateral geniculate nucleus of the thalamus and terminate in different layers of V1. L+M-, L-M+, M+L-, and M-L+ terminate in layer 4Cβ; (L+M)+S- in layer 4A and S+(L+M)- in layer 3B blobs ^9^. The neural circuits that give rise to color-opponent mechanisms and color appearance (Stage 3) require specific patterns of mixing between L/M and S/(L+M) cone-opponent mechanisms^1^ (Fig. 1) and therefore cannot occur before V1. Comparisons of visual responses between LGN and V1 indicate that these interactions likely begin within V1^10–13^. Although V1 might provide the substrates for interactions between cone-opponent mechanisms, further processing in higher visual areas, including V2, V4 and inferotemporal cortex, is likely to be required to establish the populations of neurons that encode for color perception^14–17^.

**Fig. 1.**
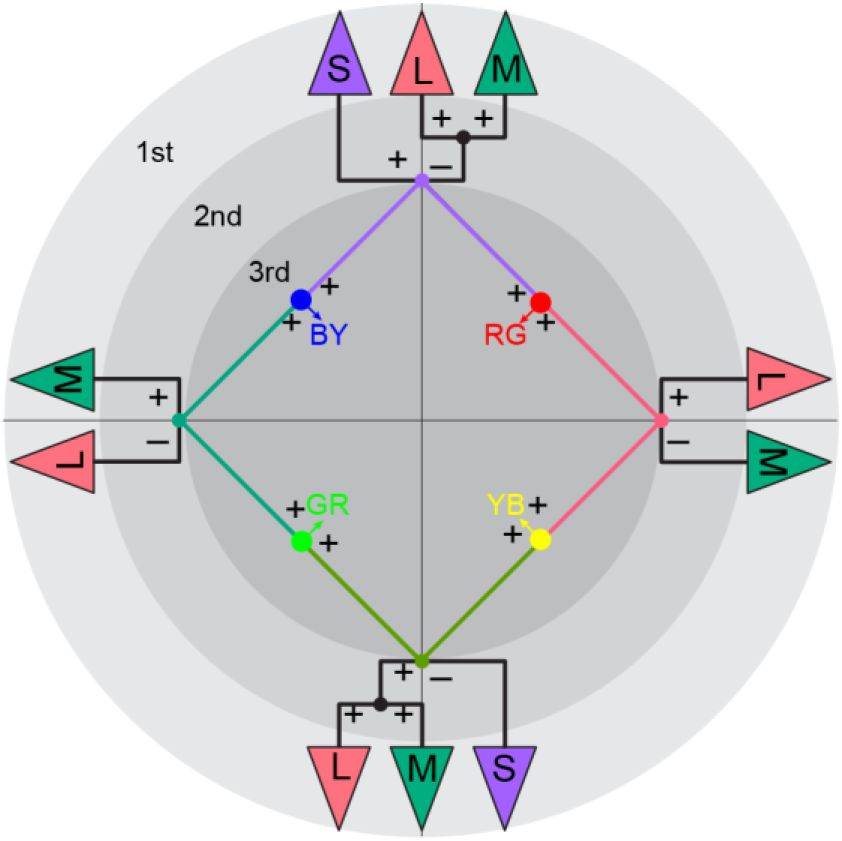
Three-stage zone model of color vision (adapted from Stockman and Brainard^1^) At the first stage (outer ring), light is transduced by three types of cone photoreceptors, L (long), M (middle), and S (short)-wavelength sensitive. In the second stage, cone signals are integrated through four cone-opponent mechanisms (L+M-, M+L-, S+(L+M)- and (L+M)+S-) instantiated by retinal circuits connecting specific cone types to retinal ganglion cell types. At the hypothesized third stage, cone-opponent signals are predicted to combine in four specific combinations to generate color-opponent mechanisms, as well as neurons whose activities underlie the perception of four “unique colors” (red – RG; green – GR; yellow – YB; blue – BY).

The expected cone-opponent mixing patterns of Stage 3 (Fig. 1) are based on explicit models and circuit level predictions generated by the psychophysics community to account for the appearance of the opponent colors, red and green or blue and yellow^18,19^, as well as “unique colors”^1,20–22^. Specifically, the appearance of each of the four “unique colors” (red – RG; green – GR; yellow – YB; blue – BY) requires a particular mixture of the possible combinations between L/M opponent and S+ or S- mechanisms, which are L+S+, M+S-, L+S-, and M+S+, respectively. Other combinations, such as L+ with M+ and S+, or L- with M- and S-, would create achromatic responses. Since both color selective and achromatic visual responses are common in V1, the full range of observed responses could theoretically be generated through random circuits that would implement all possible combinations and generate a range of color and achromatic selectivities at the level of individual neurons. Arguing against this possibility, many studies using single cell recording ^23–25^, intrinsic signal imaging (ISI) ^25–28^ and 2-photon calcium imaging (2PCI) ^29,30^ have found evidence for columnar color organization and at least some degree of separation of color-selective and achromatic neurons. The most recent of these studies reveals micro-maps of cone responses in V1 with an organization that could potentially provide a substrate for cone-opponent signal mixing and implementation of color appearance mechanisms^30^.

Here we present data showing that the predicted combinations for mixing cone-opponent mechanisms to generate color appearance mechanisms are implemented by functionally-organized circuits in the superficial layers of V1. We developed intrinsic signal imaging (ISI) methods to map the locations in V1 that are responsive to the ON and OFF phases of cone-isolating stimuli across large swaths of cortical territory. These maps reveal cone-opponent functional domains (COFDs) where neurons are preferentially activated by the ON or OFF phases of cone-isolating stimuli. Results show that L-ON (L+) and M-OFF (M-) responsive regions are always spatially overlapping, as are L-OFF (L-) and M-ON (M+) regions, to generate L+M- and M+L- domains. The L+M- and M+L- domains occur as adjacent non-overlapping pairs. S+ and S- domains are also organized as non-overlapping pairs. Importantly, the axes joining the L+M−/M+L- pairs are often organized roughly orthogonally or interdigitated in parallel to the axes joining nearby S+/S-pairs, to create interactions across all four combinations. The same relationships are observed for the underlying individual neurons assayed with 2-photon calcium imaging (2PCI) and the intersections of the interacting pairs often form hue “pinwheel” or “linear” structures. These combinations correspond to the circuits predicted to generate color-opponency and unique hues RG, GR, YB and BY in third stage mechanisms (Fig. 1). Finally, ISI hue-phase maps and 2PCI of individual neurons reveal that the functional architecture and micro-architecture of neuronal color preferences within intersecting COFDs is as expected from relatively linear combinations of these cone-opponent signals, indicating that circuits in these regions do in fact implement the predicted mixing.

## Results

We conducted ISI and 2PCI to map the functional architecture and measure the visual responses of neurons in superficial layers of V1 using methods described previously^29^. Here we present data collected from 7 adult male and female macaque monkeys (*M. fascicularis*) (see Extended Data Table 1). All data were collected from anesthetized animals following procedures approved by the Salk Institute Institutional Animal Care and Use Committee. We present ISI data from 5 animals (A1, A2, and A5 to A7) and 2PCI data from 3 animals (A1, A3, A4). A previous publication^29^ described limited data from 2 of the animals (A3, A4) subjected to 2PCI; data presented here include responses to visual stimuli and analyses not described in the previous publication.

### Cone-opponent functional domains in V1

We used the 2PCI and ISI imaging methods described above to reveal cone-opponent functional domains in V1. We begin by describing the functional micro-architecture of cone inputs to individual neurons measured with 2PCI imaging. We presented flashed sine-wave gratings (Fig. 2a) while monitoring changes in fluorescence of superficial V1 neurons (150-310 μm depth, Fig. 2b-d) expressing the genetically-encoded calcium indicator GCaMP6f (Fig. 2c, d). These sine-wave gratings consisted of 4 sets with different colors which were L, M, or S cone-isolating or achromatic, and they were presented in separate stimulus blocks. Each set was generated using the same Hartley basis function^31^, therefore, the orientations, spatial frequencies (SFs), and spatial phases were identical between each set. Fluorescence changes were converted to inferred spike-rate time sequences that were then used to calculate spike-triggered averages (STA) of stimuli as linear estimates of receptive fields (Fig. 2e-h). 2PCI from 9 imaging regions (16 planes, 1168 × 568 μm or 1100 × 725 μm) yielded data from 13117 neurons that were visually responsive to drifting gratings. A subset of those visually responsive neurons (2420 of 13117 neurons, 18.4%) had significant STA kernels to one or more of the Hartley stimulus sets. These included 1578 neurons with significant STAs to the achromatic stimulus set, 596 to L-cone isolating, 607 to M-cone, and 600 to S-cone. That only a minority of neurons have significant STA kernels is expected from the prevalence of complex cells in superficial V1 that do not linearly integrate responses to flashed grating stimuli. The smaller numbers of neurons responsive to L and M cone-isolating versus achromatic stimuli is expected from the low cone-contrasts of the L and M cone-isolating stimuli (18%-18.5%, L and M cone-contrasts were matched) imposed by the properties of the CRT monitor, versus 98%-99% achromatic stimulus contrast. The small minority of neurons with significant S-cone STA kernels cannot be attributed to the cone contrast of the S-cone stimulus, which was 89.9%-90.2%.

**Fig. 2.**
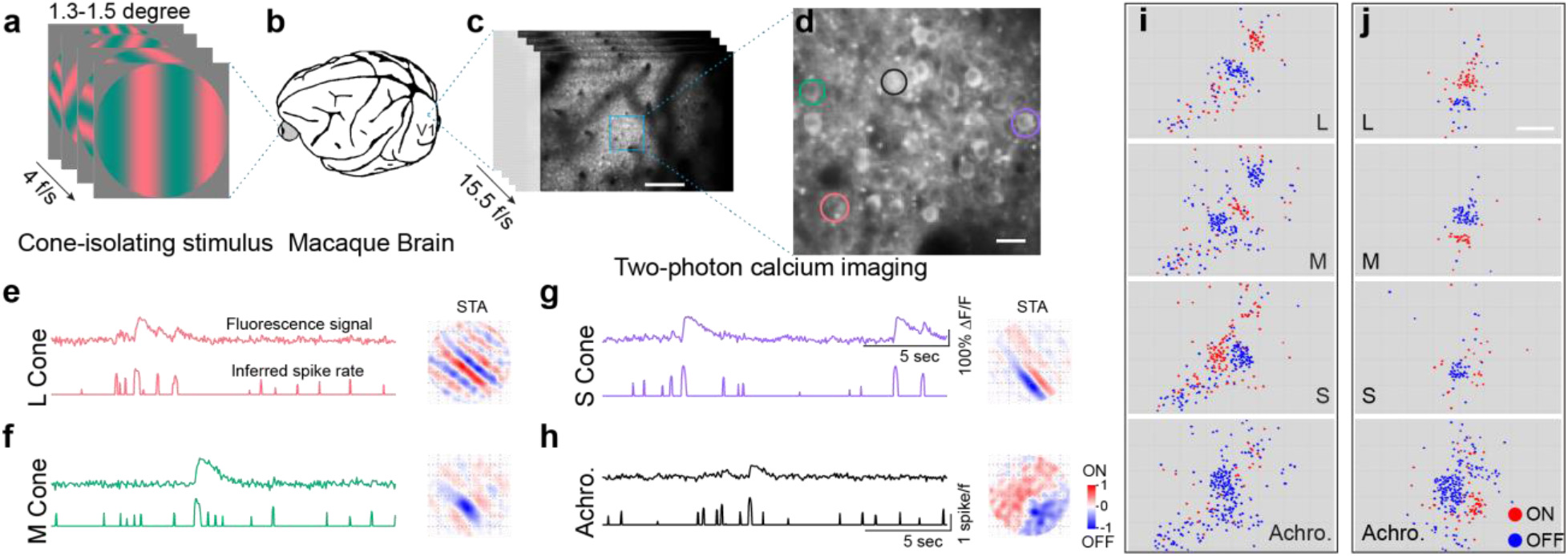
Clustering of neurons with ON/OFF-dominant receptive fields. **a-c**, Presentation of flashed sine-wave gratings (**a**), L-cone isolating is shown as an example, 4 frames per second to the anesthetized macaque monkey (**b**) during simultaneous two-photon calcium imaging (**c**) in primary visual cortex (V1). Scale bar in **c** is 200 μm. **d-h**, Higher magnification view of the imaging region (**d**, scale bar: 20 μm) with four neurons selected as examples to show their fluorescence signals, inferred spike rates, and their receptive fields computed from spike-triggered average (STA) shown in **e-h**. The color of traces in **e-h** are matched with the color of circles in **d** to indicate the origin of the signals. Responses of the four different neurons to four different stimuli, L-cone isolating (**e**), M-cone isolating (**f**), S-cone isolating (**g**), or achromatic (**h**) flashed sine-wave gratings, are shown as four panels. The fluorescence signal (upper trace in each panel) is normalized to the response to blank condition (*ΔF/F*). The spike rate (lower trace in each panel) is inferred from raw fluorescence signal. Red colors in the STA images indicate ON sub-regions of spatial receptive fields, and blue indicates OFF sub-regions. Each grid in the STA images is 0.2 degree, and total size is 1.6 degree. **i-j**, Functional maps of the locations of neurons with ON-dominant (red dots) and OFF-dominant (blue dots) receptive fields in response to each stimulus type (L, M, and S cone isolating and achromatic (Achro.)). Neurons are organized in ON and OFF clusters. **i-j** are two imaging regions from animal A1 (more examples are shown in Extended Data Fig. 1); both panels illustrate results from two merged imaging planes. Scale bar in **j**: 200 μm; applies to all panels in **i** and **j**.

As expected from previous studies mapping receptive fields of V1 neurons in response to cone-isolating stimuli^32–34^, receptive field structures varied between stimulus sets and between neurons. We do not further consider the fine spatial structure of receptive fields here. If a neuron had a significant STA kernel to one or more of the flashed sine-wave gratings sets, then we calculated its ON or OFF dominance (see Methods), and plotted the ON/OFF dominance map in red/blue, based on the locations of neurons. Maps from all imaging regions are presented in the main and supplementary Figures. (Fig. 2-3 and Extended Data Fig. 1). Here, we call attention to representative examples from two imaging regions in Fig. 2i,j and Fig. 3f-i. In all imaging regions, neurons with significant STA kernels occupied a small and relatively contiguous region within the imaging window. This organization is best appreciated in the context of maps showing the locations of all visually responsive neurons, relative to the locations of neurons with significant STA kernels (compare Fig. 5f-i to Fig. 3f-i). It is apparent from these maps that neurons with dominant responses to the ON versus OFF phases of each stimulus set are clustered together for each of the four stimulus sets. Furthermore, clusters of neurons with L-ON dominance overlap with M-OFF dominance clusters, while L-OFF clusters overlap with M-ON. The smaller numbers of neurons with significant S-cone STA kernels prevents definitive identification of the spatial relationships of S-ON and S-OFF clusters to the L and M clusters based on this relatively subjective assessment. (We present detailed neuron-by-neuron quantitative comparisons of the signs and magnitudes of cone weights to each cone-isolating stimulus below, Fig. 5m). These cone and achromatic ON/OFF dominance maps suggest that within V1 there are functional architectures where neurons with significant STA kernels are clustered together, that ON and OFF dominant neurons of each type are further clustered, and that the organized overlap between L and M cone clusters creates L+M- and M+L- cone-opponent clusters.

**Fig. 3.**
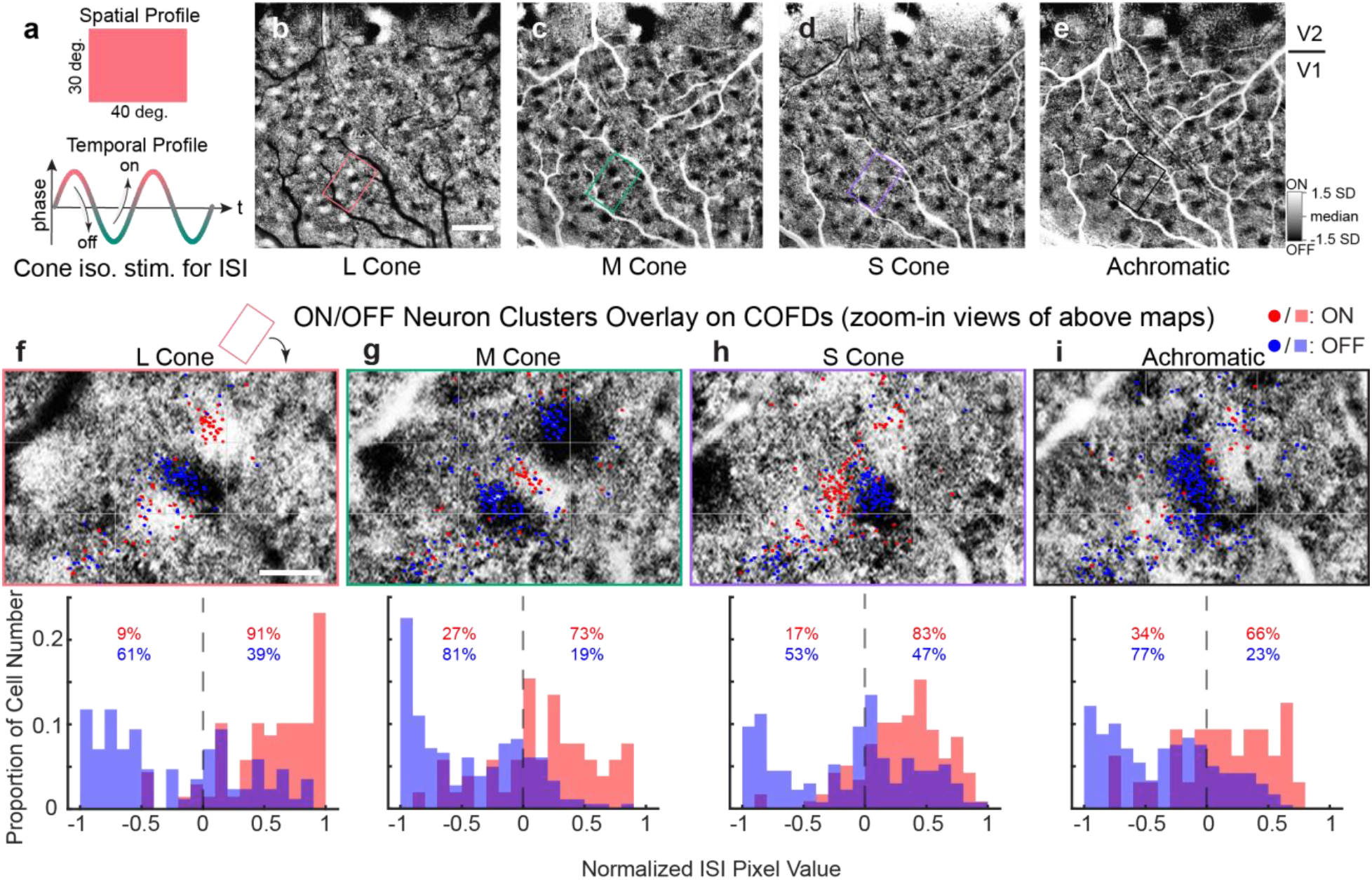
COFDs obtained from ISI are aligned with ON and OFF neuron clusters from 2PCI. **a**, Continuous periodic full-field cone-isolating and achromatic stimuli (top) with corresponding sinusoidal temporal modulation (bottom, L-cone isolating stimulus is shown). **b-e**, ISI phase maps reveal ON and OFF functional domains resulting from L-, M-, S-cone isolating and achromatic stimuli, respectively. Scale bar: 1mm. **f-i**, ON/OFF neuron clusters aligned with COFD. Panels at top: 2PCI responses of neurons (colored dots) to L-, M-, and S-cone isolating and achromatic stimuli superimposed on corresponding aligned zoom-in ISI phase maps from regions outlined in **b-e**. Individual neurons imaged with 2PCI depicted based on whether STA receptive field was predominantly ON (red) or OFF (blue). Histograms at bottom: The ON (red) and OFF (blue) neuron distributions according to the normalized pixel values of aligned ISI phase maps. Positive pixel values represent ON domains of COFDs, negative values represent OFF domains of COFDs. Numbers in the panels are the percentages of ON neurons (red text) or OFF neurons (blue text) in ON versus OFF domains. Scale bar in **f**: 200 μm; applies to **f-i**. Note that grids were added in **f-i** as landmarks for comparison. The red rectangle above **f** indicates how selected region in **b** is rotated to display **f**; same operation applies to **g-i**.

We inferred that this cone-opponent functional architecture might be visualized across much larger V1 regions using ISI, and allow its relationship to cytochrome oxidase (CO) “blobs” and “interblobs”^35^ to be revealed by aligning postmortem CO histology to ISI maps^29^. To generate ISI ON/OFF phase maps corresponding to the ON/OFF phases of L, M and S cone-isolating and achromatic stimuli, we presented full-field stimuli that were temporally modulated between ON and OFF phases of each stimulus type (L, M, S, and achromatic) at 0.017 Hz, 0.08Hz or 0.1 Hz (Fig. 3a). Subsequent Fourier analysis of the ISI images yielded ON/OFF phase maps for each stimulus type^36^. In all 5 animals imaged, and in response to each stimulus type, the phase maps showed clear patches of ON and OFF signal surrounded by regions with no phasic response, extending across the full extent of V1 (Fig. 3, 4, 6, Extended Data Fig. 1, 3b). (Phase maps in the adjacent visual cortical area V2 are not further explored here.) Furthermore, within each patch, there were invariably adjacent pairs of alternating ON and OFF domains. Because of the unknown latency of hemodynamic signal, correct signs cannot be assigned to the phase maps without additional measurements from another modality with greater temporal sensitivity. We identified the correct phases using electrophysiological recordings (Extended Data Fig. 9) within imaged domains for 2 of 5 animals (A2, A5) and using 2PCI for 2 of the remaining animals (A1, A6). For the last animal (A7), similar phase maps were generated (Extended Data Fig. 2b), but we did not identify the phase signs. We assume that the hemodynamic signal latency is consistent between every sub-region within V1 for each animal. Alignment of ISI phase maps to the ON/OFF dominance map measured by 2PCI demonstrates a clear correspondence between the two imaging modalities (Fig. 3f-i and Extended Data Fig. 1). We refer to the V1 regions with strong ON and OFF phasic responses to temporally modulated cone-isolating stimuli as cone-opponent functional domains (COFDs).

**Fig. 4.**
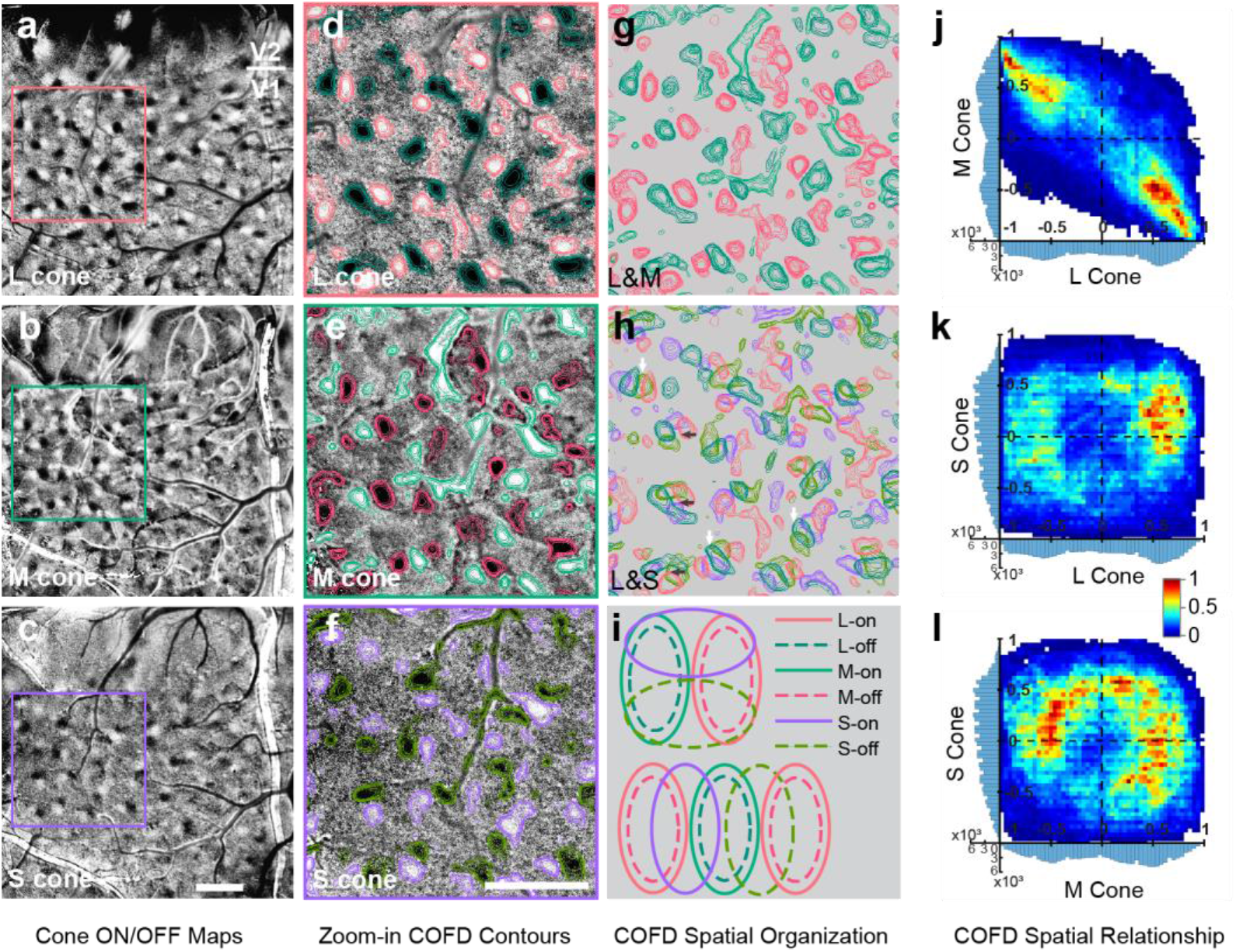
Spatial relationships between L-, M-, and S-COFDs. **a-c**, COFDs from ISI phase maps in response to full-field periodic L-, M-, and S-cone isolating stimuli, respectively. Scale bar in **c**: 1 mm; applies to **a-c**. **d-f**, Zoom-in views of maps in **a-c**. Color contours for ON and OFF domains are overlaid on images based on ISI pixel values. Scale bar in **f**: 1 mm; applies to **d-h**. **g-h**, Merge L-COFD contours (**d**) with M-COFD contours (**e**) and merge L-COFDs (**d**) with S-COFDs (**f**) to demonstrate spatial relationships between L-, M-, and S-COFDs. Same scale as **d-f**. Additional examples are shown in Extended Data Fig. 2a. L- and M-COFDs overlap extensively with an anti-phase relationship; L-on COFDs align with M-off COFDs, and L-off align with M-on (**g**, also see **j**). In contrast, S-on and S-off COFDs tend to fall between and partially overlap with L+M- and M+L-COFDs (**h**, also see **k**, **l**) with a cross-overlap organization. **i**, Schematic diagrams of the COFD relationships often seen in **h**. L+M−/M+L-pairs are often grouped with an S+/S-pair such that the pairs have orthogonal axes (**h**, dark arrows, schematized at top of **i**). Alternatively, the pairs are arranged in parallel with S+ and S-COFDs interdigitating between L+M- and M+L-COFDs (**h**, white arrows, schematized at bottom of **i**). **j-l**, 2D histograms of the spatial relationship between L, M, and S COFDs. The 1D histograms on the *x* and *y* axes display the distribution of normalized pixel values in each category. (Additional cases are shown in Extended Data Fig. 2a). The anti-phase relationships between L and M COFDs are apparent in **j**, while interactions between L or M with S-COFDs have more balanced interactions across all phases (**k** and **l**).

**Fig. 5.**
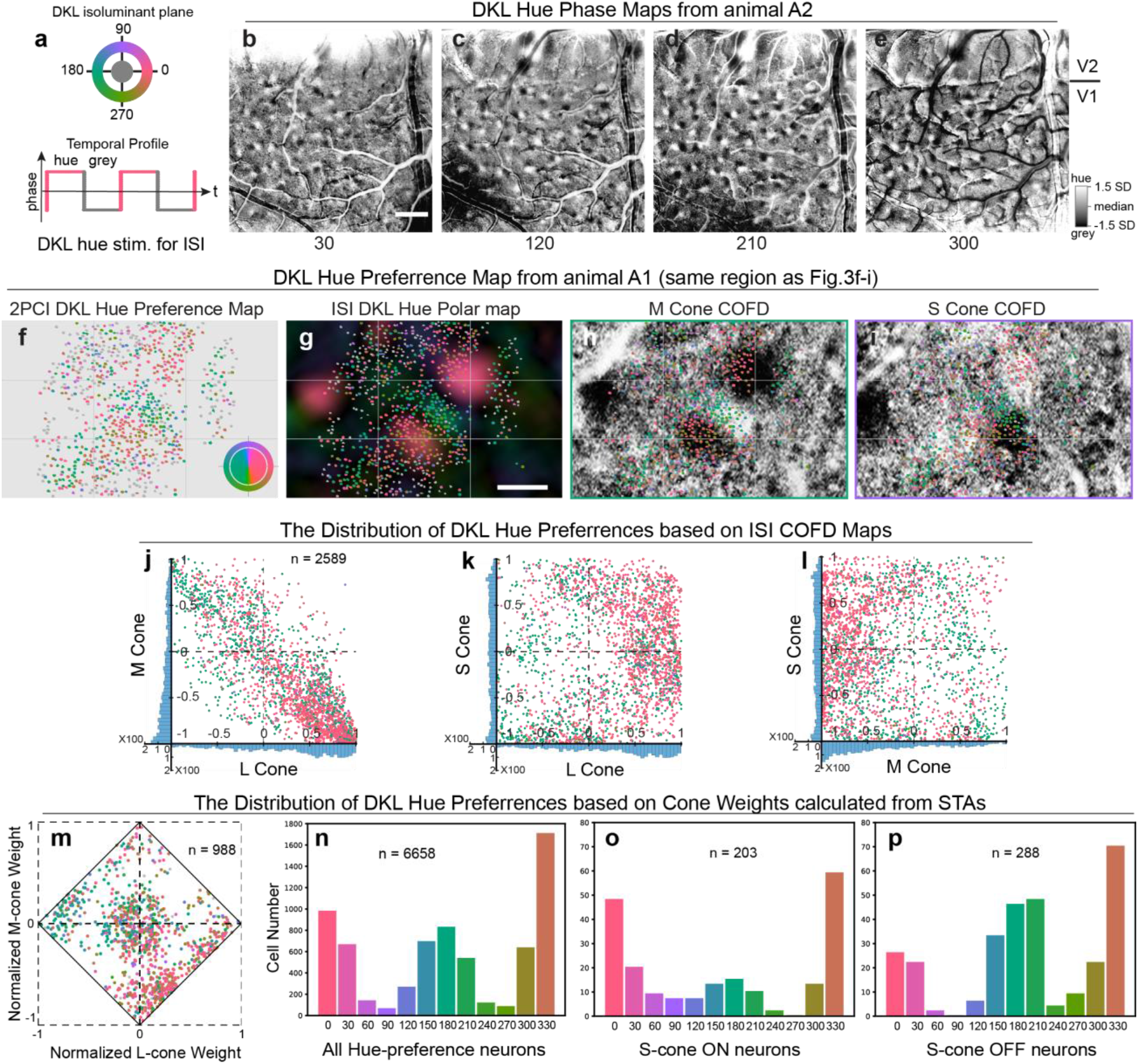
DKL hue map and its relationships to COFDs and Cone weights. **a**, Schematic of stimulus presentation. During ISI, twelve evenly spaced DKL hues were presented as full-field, square wave temporally modulated stimuli on a grey background. L+/M-, L−/M+, S+, and S-directions correspond to 0, 180, 90, and 270 degree hue directions, respectively. **b-e**, ISI hue-phase maps in response to 30, 120, 210, and 300 degree hue directions (animal A2, other maps and another case, A1, are shown in Extended Data Fig. 4a). Maps have ON/OFF (hue/grey) phase pairs. Scale bar in **b**: 1 mm; applies to **b-e**. **f**, DKL hue preference map from the same imaging region as shown in Fig. 3f-i. Individual neurons are depicted based on the DKL hue to which they responded most strongly. Neurons that were not hue selective are plotted in grey. The inserted color key indicates the relationship between the hues for plotting the map (outer ring, DKL hues have maximal contrast that can be generated by our CRT monitor) and the hues used for stimulation (inner disk, DKL hues have matched cone-increment). **g**, DKL hue preference map (the same image shown in **f**) from 2PCI overlaid on DKL hue polar map of the same region from ISI. More cases are shown in Extended Data Fig. 5. **h-i**, DKL hue preference map (the same image shown in **f**) overlaid on M-cone and S-cone COFDs (the same regions shown in Fig. 3g-h). Scale bar in **g**: 200 μm; applies to **f-i**. Note that grids were added in **f-i** for comparison purpose. **j-l**, Relationships between ISI COFD organization and DKL hue preferences based on individual neurons identified by 2PCI. Each dot in the scatter plots represents a neuron (2589 neurons) which is within COFD, and its *x y* coordinates are the mean normalized pixel values of COFD maps within that neuron (e.g., after aligned 2PCI image with ISI image, the same neuron in **h** and **i** occupies the same group of ISI pixels. see Method). Histograms on *x* and *y* axes demonstrate the distribution of neurons based on normalized pixel values from ISI. Individual neurons are colored according to their preferred DKL hue based on 2PCI. **m**, Relationships between DKL hue preferences and cone weights calculated from STAs. 988 neurons with significant L, M, or S-cone STA kernels from 5 imaging regions (10 planes, see Extended Data Fig. 5) of A1 were selected to calculate normalized cone weights. Note that not all DKL hue selective neurons have significant STA kernels. L and M cone weights are plotted on the *x* and *y* axes in the diamond plot. Due to normalization summing total absolute values of weights to 1, S-cone weights are implicit as the distance from the edges of the diamond. Each neuron in the plot is colored according to its preferred DKL hue. Neurons with significant STAs, but which are not hue selective, are plotted as grey. **n-p**, The distribution of preferred DKL hues of all hue-selective neurons (**n**), hue-selective neurons with significant S-ON STA (**o**), and hue-selective neurons with significant S-OFF STA (**p**). The preferred hue is shown as the color of bar, and the number of neurons sampled is shown in each panel. (5 imaging regions, 10 planes, from A1.)

**Fig. 6.**
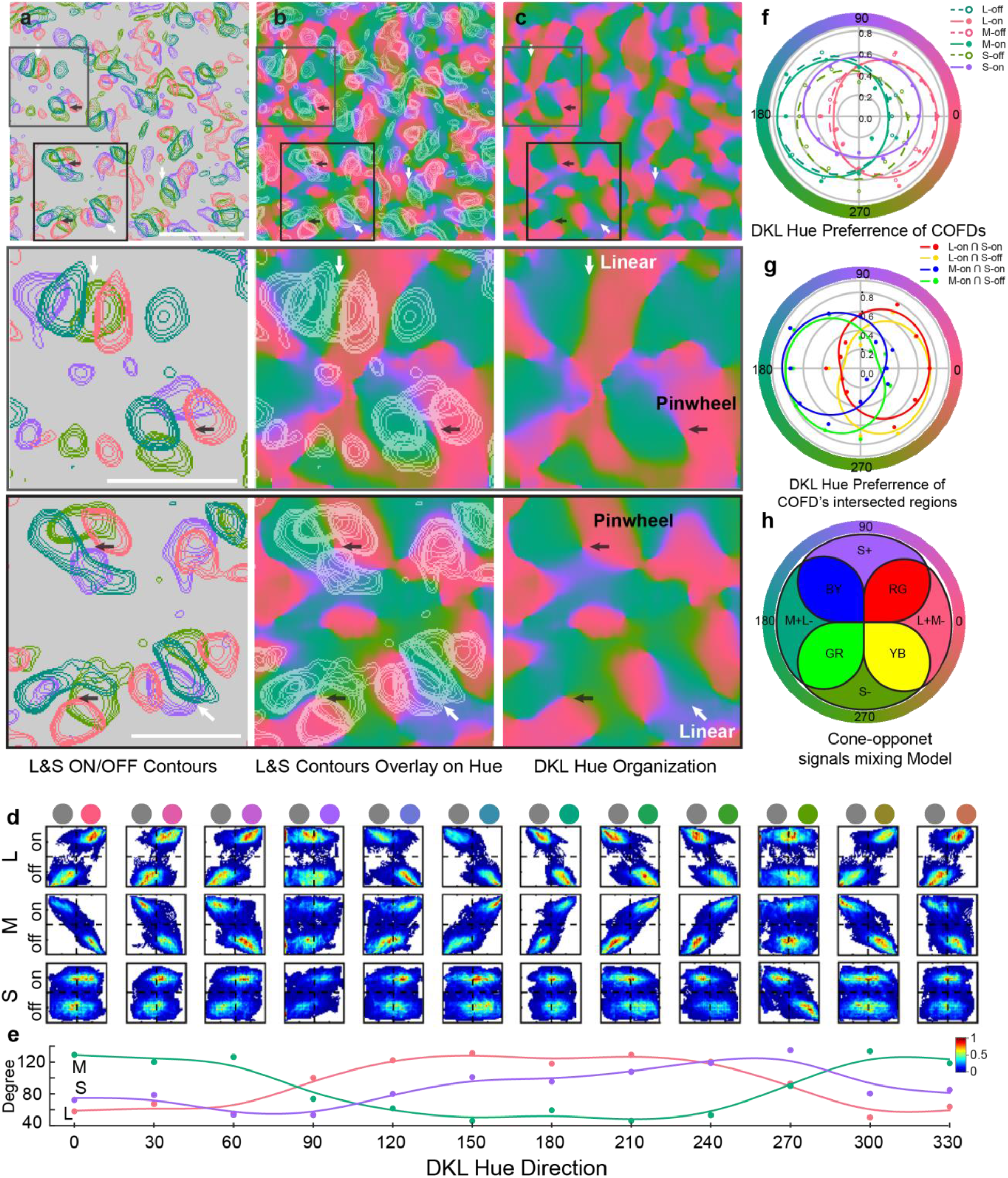
The functional organization of DKL hue direction map and its relationship with COFDs. **a-c**, COFDs contours (**a**), DKL hue direction map (**c**), and their spatial overlay (**b**). The color in DKL hue direction map is the preferred hue calculated from DKL hue phase map using vector summation, see Methods for details. Panels in the second and third rows are the zoom-in views of selected regions shown in the first row. Both pinwheel (pointed by black arrows) and linear (pointed by white arrows) functional structures exist in DKL hue direction map, and they spatially correspond to the functional organization of COFDs. Scale bar in top row of **a** is 1mm, and it is 0.5mm in the middle and bottom row of **a**; applies to **a-c**. Note that the first image in **a** is the same image in Fig. 4h. The cortical region shown in the first row of **a-c** is the same region shown Fig. 4a-h. **d-e**, Quantitative analyses of the spatial relationship between DKL hue direction domains and each type of COFD (another case, A1, is shown in Extended Data Fig. 4b,c). Rows in **d** correspond to comparisons of hue maps with L (top row), M (middle row), and S (bottom row) COFDs. Columns in (**d** and **e**) correspond to relationships to hue-phase maps generated with stimuli modulated in the hue directions indicated at bottom of **e**. **d**, The ON and OFF phases of cone-isolating (L, M, and S) stimuli are shown as ON and OFF colors of each cone type on the left side. The colors of twelve DKL hues and grey are shown as color disks on the top of the panel. The bivariate histograms show systematic relationship between COFDs and DKL hue direction domains. **e**, The angles defined by the top 10% of pixel densities in each bivariate histogram for each cone type and in relation to each hue direction are plotted. X-axis is the stimulus hue direction in DKL isoluminant plane, Y-axis is the angle calculated from each bivariate histogram. Note that the L-cone and M-cone fitted curves have a counter-phase relationship, 180 degrees apart. The phase of the S-cone fitted curve is 90 degrees from and halfway between the L and M curves. **f**, Hue tuning curves calculated from the means of ISI pixel values within each COFD region in response to each DKL hue direction. **g**, Hue tuning curves calculated from the means of ISI pixel values within each COFD-intersection region in response to each DKL hue direction. Method of curve fitting used in **f** and **g** is described in Methods. Another case is shown in Extended Data Fig. 4e,f. **h**, Cone-opponent signals mixing model based on COFDs. The spatial organization and overlap of COFDs follows Stage 3 mixing rules for color opponency and color appearance mechanisms, and creates hue tuning. The organization allows four intersections between L/M and S/(L+M) cone-opponent mechanisms at the four overlapping regions of COFDs, to generates color appearance mechanisms and encompasses the four “unique colors” (red – RG; green – GR; yellow – YB; blue – BY).

### COFD interactions follow cone-opponent signals mixing rules

Based in their spatial overlaps, the COFDs in V1 appear to follow the cone-opponent mixing rules predicted by the Three-stage model (Fig.1). In particular, L-ON functional domains overlap with M-OFF and L-OFF overlap with M-ON, but there is minimal overlap between L-ON and L-OFF or between M-ON and M-OFF (Fig. 4a, b, d, e, and g). This overlap generates L+M- and M+L- domains that appear to intersect with both S+ and S- domains (Fig. 4c, f, h). Two different types of intersections appear to be common within these maps. In many cases, an isolated set of COFDs includes one L+M−/M+L- pair and one S+/S- pair, in which the axes connecting the L/M COFDs are perpendicular to the axes connecting the S COFDs (e.g., *dark arrows* in Fig. 4h, schematic in Fig. 4i, *top*). Other interactions occur through a parallel arrangement, with S+ and S- domains interdigitated between the L+M- and M+L- domains (e.g., *bright arrows* in Fig. 4h, schematic in Fig. 4i, *bottom*). These spatial relationships potentially allow the S+ and S- domains to mix with the L+M- and M+L- domains in the combinations predicted by Three-stage model (Fig. 1 and 4i), suggesting that the COFDs are the substrate for implementation of the cone-opponent signal mixing rules that begin to generate color appearance mechanisms.

To more rigorously test this possibility, we next quantified the relationships between the L, M, and S-COFDs. The COFD map of each cone type was thresholded to generate a mask to select ON- and OFF-phase regions, then masks from all three cone type maps were merged to create a final COFD mask including L, M and S-COFD regions. Then using normalized pixel values from L, M, and S-COFD maps (range: [−1 to 1]), each pixel within the COFD mask was assigned three values corresponding to the modulation (including magnitude and sign) by each of the three cone-isolating stimuli at that pixel. Pixel-by-pixel comparisons were then made across each of the three possible comparisons of cone-types (M to L, S to L, and S to M). Typical results are illustrated in Fig. 4j-l. Similar results were observed for all 4 animals with known ON/OFF phases (Fig. 4 and Extended Data Fig. 2a). It can be seen that for the L-cone versus M-cone comparisons, the great majority of pixels (84.2%) fall into the LM opponent quadrants (L+M- and M+L-) and not in the achromatic quadrants (L+M+ and L-M-). In contrast, comparisons between S-cone and either L- or M-cone maps generate pixels relatively evenly distributed across all 4 quadrants, as predicted.

In addition to interactions between cone-opponent mechanisms within a 2-dimensional isoluminant plane (as illustrated in Fig. 1), a full accounting of color space would also include color directions that modulate along the achromatic axis. As described above (Fig. 3i), we also observe ON/OFF phase maps in response to full-field achromatic stimuli; these overlap extensively with and have a similar structure to COFDs (Fig. 3; Extended Data Fig. 3). To assess whether the achromatic ON/OFF domains might contribute to generation of a diversity of selectivities for color directions with achromatic modulation, we quantified the relationships between COFDs and achromatic domains, as described above for the relationships between the three COFDs. Pixel-by-pixel comparisons between the achromatic domains and each of the COFDs (L, M, or S) are illustrated in Extended Data Figs. 3d-f. Consistent with a possible contribution of achromatic modulation to color tuning of the underlying neurons, both the ON and OFF achromatic domains overlap extensively with both ON and OFF regions of all three types of COFDs. This contrasts with an alternate hypothesis in which the achromatic domains do not contribute to color tuning and would instead only overlap with COFD pixels of matching sign. Such same-sign overlap would be expected to generate relatively achromatic responses without substantial chromatic modulation.

### DKL hue direction maps and hue pinwheels

The above observations show that the overlap of the COFDs follows the rules required to integrate cone-opponent signals in the combinations that would generate color opponency, but do not assess functional mapping that might be present to a larger range of intermediate colors predicted from mixing within COFDs. They also do not test whether mixing occurs at the level of individual neurons or how neurons within the overlapping regions respond to a larger range of hues. For example, are the ISI phase responses to colors that concurrently modulate multiple cone types predictable from the COFD maps, as expected from mixing of cone-opponent inputs? Do the individual neurons within the COFDs prefer colors that would be predicted from mixing of the cone-opponent signals within the COFDs?

To address these questions, we selected as visual stimuli a set of 12 hues spaced evenly in the isoluminant plane of DKL color space^37^. We chose the DKL isoluminant plane because there is a close correspondence between DKL hues and the cone-opponent signals^10^, allowing straightforward predictions of hue preferences expected from linear integration of cone-opponent inputs. The 12 DKL hues were presented as drifting gratings for 2PCI or temporally modulated full-field stimuli for ISI, and imaged the responses of neurons within the COFDs. We generated both ISI hue-phase maps (Fig. 5b-e and Extended Data Fig. 4a), and 2PCI hue maps (Fig. 5f,g and Extended Data Fig. 5) to reveal the overall functional architecture and the microarchitecture of hue preferences of individual neurons. Within the DKL isoluminant plane, hue directions of 0 and 180 deg. correspond to the L+M- and M+L- directions and appear pinkish and greenish, respectively, while 90 and 270 deg. correspond to S+ and S- and appear violet and lime (Fig. 5a and Extended Data Fig. 8b). Intermediate hues are predicted from mixing of the cone-opponent signals that modulate along these cardinal axes. The L, M, and S cone increments were matched for the DKL stimuli presented to animal A1 (both ISI and 2PCI experiments), while for animal A2 (ISI experiments only) all cone increments were set to the maximum cone contrasts achievable with the monitor, substantially increasing the S-cone contrasts relative to L and M cone contrasts (Extended Data Fig. 8b). These differences are noted where relevant, below.

Similar to ISI phase maps generated with cone-isolating stimuli (COFDs), the hue-phase maps generated using DKL stimuli were composed of ON/OFF pairs with strong signals surrounded by regions with weak or no phasic modulation (Figs 5a-e). We used these data to generate DKL hue direction maps (Figs. 6b,c and Extended Data Fig. 4d) using methods similar to those typically used to calculate orientation maps^38,39^; the hue-phase-regions from 12 hue-phase maps were selected, vectorized and summed pixel-by-pixel to generate the hue direction map. We then scaled the hue direction map with the vectors’ magnitude to generate hue polar maps (Figs. 5g and Extended Data Fig. 5). The resulting hue direction maps have a surprising and striking resemblance to orientation maps^38,40–43^ and motion direction maps^39,44,45^, including pinwheel-like structures (Fig. 6a-c, arrows highlight selected pinwheel centers) surrounded by linear iso-hue direction regions. (See also orientation and hue polar maps in Extended Data Fig. 5). In some striking cases the pinwheel centers correspond precisely to regions of convergence of L+M−/M+L- COFD pairs with S+/S- pairs whose axes are arranged orthogonally (Fig. 6a-c, *dark arrows*; note that Fig. 6b is generated by overlaying contours in Fig. 6a on top of Fig. 6c), but other pinwheel centers are found where COFD pair axes interact in a more parallel arrangement (Fig. 6a-c, *white arrows*), and some are even found outside the COFDs. DKL hue pinwheels were most apparent when generated using stimuli with cone increments maximized (Fig. 6a-c), but were also observed when cone increments were matched (Extended Data Fig. 4d).

### COFDs and hue preferences of individual neurons

We next assessed whether individual neurons within the COFDs prefer colors that would be predicted from mixing of the cone-opponent signals within the COFDs. Figs. 5f-g illustrate, for one imaging region (out of 5 sampled), the 2PCI-based functional microarchitecture for hue preferences in the DKL isoluminant plane (matched cone increments) and the overlaps with COFD maps are shown in Fig. 5h-i. (Maps from all 5 imaging regions, 10 planes, are shown in Extended Data Fig. 5). All visually responsive neurons are indicated, with significantly hue-selective neurons identified by their preferred hue and non-selective neurons in grey. It is readily apparent that neurons with similar hue preferences are grouped together and that hue preferences shift gradually across the cortical surface (Fig. 5g and Extended Data Fig. 5). When overlaid on the M-cone COFDs, it is apparent that neurons in the M+ (L-) domains prefer greenish hues while those in the M- (L+) domains prefer pinkish hues (Fig. 5h). The predominant red and greenish preferring neurons are both found in both S+ and S-domains (Fig. 5i).

To quantitatively evaluate the relationships between the preferred DKL hues of individual neurons and their locations within ISI COFD maps, we first selected neurons located within COFDs and that were significantly tuned to DKL hues (*n* = 2589 neurons from 5 imaging regions). We then generated a mask for each neuron to identify its corresponding ISI pixels and the normalized pixel values from L, M, and S-COFD maps (see above) for each neuron were averaged to yield the magnitudes and signs modulated by L, M and S cone-isolating stimuli. Similar to Fig. 4j-l, three possible comparisons of cone-types (M to L, S to L and S to M) were plotted neuron-by-neuron (Fig. 5j-l), and each neuron was colored according to its preferred DKL hue measured by 2PCI. For the L cone versus M cone plot (Fig. 5j) pixels from the great majority of neurons (87.1%) are in the L/M opponent quadrants; neurons with pixels in the L+M- quadrant are significantly more likely to prefer DKL hues near 0 deg. (0 ± 60 deg., 1167 of 1650 neurons, 70.7%) than near 180 deg. (180 ± 60 deg., 462 of 1650 neurons, 28.0%; *P* < 0.0001, Fisher’s exact test); and neurons with pixels in the M+L- quadrant are significantly more likely to prefer hues around 180 deg. (180 ± 60 deg., 344 of 605 neurons, 56.9%) than around 0 deg. (0 ± 60 deg., 250 of 605 neurons, 41.3%; *P* = 0.0018, Fisher’s exact test). For the S-cone versus L and M cone comparisons (Fig. 5k, l) neurons are more evenly distributed amongst the 4 quadrants, as expected from the overlap of both S+ and S- domains with all combinations of L and M cone domains.

Perceived color is influenced not only by the relative activation of different cone types, but also by luminance, or achromatic contrast. For example, the DKL color space (like other color spaces) includes modulation not only in the isoluminant plane, as we have explored here, but also along the achromatic axis^37^. As described above, we have observed achromatic ON and OFF ISI phase maps similar to the COFDs (Figs. 2i,j and 3e,i), and their overlap with COFDs suggests a possible role in the generation of hue preferences (Extended Data Fig. 3). But we have not explored the responses of individual neurons to the much larger 3D hue stimulus space. Future studies should explore how the achromatic ON and OFF domains might interact with the COFDs to generate predictable tuning across the full 3D color space.

### COFDs and DKL hue direction maps

Consistent with 2PCI results indicating mixing of cone-opponent signals to generate intermediate hue-selectivities of individual neurons within COFDs (above), we also observed gradual shifts in ISI hue-phase maps (Fig. 6d,e and Extended Data Fig. 4a-c) generated with stimuli modulated at non-cardinal directions in the DKL isoluminant plane (Fig. 5a). These maps were generated using DKL stimuli with cone increments maximized (non-matched, see above) to better reveal the locations with responsiveness to S-cones. The gradual shifts in ISI hue-phase maps are quantified in Figs. 6d and 6e. Similar to comparisons between normalized ISI phase pixel values and signs across maps from the cone isolating stimuli (Fig. 4j-l), each row of Fig. 6d illustrates comparisons between pixel values from each of the 12 hue-phase maps with either the L (top row), M (middle row) or S cone (bottom row) phase maps. It can be seen that as DKL hue directions shift, the phase relationships to the COFD phases shift gradually (Fig. 6d,e and Extended Data Fig. 4b,c). When hue directions are near the L/M axis (0 and 180 deg.) pixels are in opposing quadrants in relationship to the L and M phase maps (Fig. 6d and Extended Data Fig. 4b) and accordingly the angles joining the two clusters of pixels in each plot are near 45 and 135 deg. (Fig. 6e and Extended Data Fig. 4c). When these hue phase maps (0 and 180 deg. hues) are related to the S-cone phase maps, the pixels occupy all 4 quadrants (Fig. 6d and Extended Data Fig. 4b) and joining angles are near 90 deg. (Fig. 6d and Extended Data Fig. 4c). As hue directions gradually increase in S cone modulation, comparisons to the L and M phase maps show pixels gradually occupying all 4 quadrants (Fig. 6d and Extended Data Fig. 4b) and joining angles gradually shifting (Fig. 6e and Extended Data Fig. 4c), until at maximal S cone modulation (90 and 270 deg) all 4 quadrants are occupied and joining angles are near 90 deg. In contrast, comparisons of hue phase maps to the S-cone maps show pixels more evenly distributed across all 4 quadrants, with the exception of comparisons to the S+ and S- hues (90 and 270 deg.) where opposing quadrants are occupied (Fig. 6d and Extended Data Fig. 4b) and joining angles are near 45 and 135 deg. (Fig. 6e and Extended Data Fig. 4c). These observations indicate that, not only does the overlap between COFDs follow the rules predicted to instantiate color appearance mechanisms (Fig. 1 and 4j-l), but that the interactions between L, M, and S-COFDs generate preferential responses to intermediate hues within the DKL color space, as predicted from those interactions (Figs. 5j-l, 6d-e, and Extended Data Fig. 4b-c).

### Cone inputs and hue tuning

For 2PCI-assayed neurons with significant STA kernels to the L, M, or S cone-isolating stimulus set, their preferred hues in color space were also highly consistent with the magnitude and sign of cone inputs estimated from the STA kernels. This was the case both for hues presented in the DKL isoluminant plane (Fig. 5m) and in CIE color space^29^ (Extended Data Fig. 6a). This relationship is most apparent from the responses to DKL hues, due to the clear correspondence between cone-opponency and DKL hue directions. Normalized cone weight plots for both DKL and CIE hues (Fig. 5m and Extended Data Fig. 6a) show that the great majority of neurons are in the L+M- and M+L- cone-opponent quadrants, as expected from the mixing described above. Neurons in the achromatic quadrants have only weak STAs to one or another of the L or M cone-isolating stimuli, as expected from the noise inherent to calculation of input magnitude from STA kernels and the methods we used to assign values. (Neurons with a significant STA to only one stimulus set were nevertheless assigned values for the non-significant stimuli rather than pinning values to zero.) As expected, for the neurons tested with DKL hues, the great majority of hue-selective neurons in the L+M- quadrant preferred DKL directions around 0 deg. (315 of 390 = 80.8% of neurons at 0 ± 60 deg.), while those in the M+L- quadrant preferred DKL directions around 180 deg. (119 of 213 = 55.9% of neurons at 180 ± 60 deg.). For neurons tested with CIE hues, hue-selective neurons in the M+L- quadrant nearly all preferred blue while neurons in the L+M- quadrant mostly preferred red with some preferring blue (Extended Data Fig. 6a). This arrangement is expected from the stronger L cone contrast for red and stronger M cone contrast for blue (Extended Data Fig. 8c), combined with the fact that the OFF phase of the blue stimulus can sometimes generate a stronger L response than M. It is important to note that hue selective neurons generate both ON and OFF responses to drifting hue gratings and due to the slow dynamics of calcium signals observed with 2PCI, it is not possible to definitively identify whether responses are being generated to the ON versus the OFF phases of the drifting gratings. While ON responses are generally stronger than OFF responses, this may not always be the case. Therefore, the hue preferences observed from 2PCI imaging of responses to drifting gratings are not expected to always match the predictions from responses to flashed, cone-isolating stimuli.

There is also a clear separation of a set of neurons near the middle of the diamond plots with relatively strong S-cone STA kernels (Fig. 5m and Extended Data Fig. 6a). Note that the S-cone isolating stimulus used to generate these STA kernels is much higher contrast (89.9%-90.2%) than the L and M-cone stimuli (18%-18.5%), contributing strongly to the separation of this group. This is apparent from the overall distribution of hue preferences to DKL stimuli (Fig. 5n). Here it can be seen that when stimuli are matched for cone increment magnitude, responses are dominated by colors modulated at or near the L-M opponent axis (0 and 180 deg.). Extremely few (138 of 6658 neurons, 2.07%) preferred stimuli modulated in the S+ or S- directions. Nevertheless, S-cone inputs could be seen to have a significant, although relatively weak influence on preferred DKL hue. Amongst neurons with significant S-cone STA kernels, their preferred directions in the DKL isoluminant plane were consistent with expected shifts resulting from linear integration with L/M opponent signals. Neurons with significant S-cone ON STA kernels were more frequently tuned to the DKL hue at 90 deg. than neurons lacking significant S-cone STA kernels (7 of 203 neurons, 3.45% versus 52 of 6167 neurons, 0.84%; *P* = 0.0058, Fisher’s exact test), or than neurons with significant S-cone OFF STA kernels (0 of 288 neurons, *P* = 0.0045, Fisher’s exact test). Similarly, neurons with significant S-cone OFF STA kernels were more likely to prefer the DKL hue at 270 deg. than neurons without significant S-cone kernels (9 of 288 neurons, 3.13% versus 70 of 6167 neurons, 1.14%; *P* = 0.0184, Fisher’s exact test), or than neurons with significant S-cone ON STA kernels (0 of 203 neurons, *P* = 0.0175, Fisher’s exact test).

### Preferred hues within COFDs and their intersections

The relatively weaker influence of the S cone relative to the L/M opponent signals during interactions between the COFDs can be seen not only in the hue tuning of the individual neurons (see above), but also in the tuning of ISI pixels within the different COFDs to the DKL stimuli. To further quantify the relationships between COFDs and DKL hue responses, the COFD masks were used to identify pixels in each COFD ON or OFF region (Fig. 6f and Extended Data Fig. 4e), as well as intersections between regions (Fig. 6g and Extended Data Fig. 4f), and then the ISI hue responses of those pixels were used to generate hue tuning curves. For example, it can be seen in Fig. 6f that, as expected, the pixels in the L+ and M-COFDs were most strongly modulated by hues at or near the 0 deg. hue direction and most weakly modulated by the 180 deg. hue direction. In contrast, the M+ and L- pixels were most strongly modulated at 180 deg. and least strongly at 0 deg. Importantly these tuning curves are relatively symmetric around the 90 to 270 deg. axis, as expected from near equal influences of S-ON and S-OFF signals, resulting from the symmetric mixing of S-ON and S-OFF COFDs with the L+M- and M+L- domains. Also as expected, the S+ COFDs responded more strongly to 90 deg. than to 270 deg. and S- more strongly to 270 deg. than to the 90 deg. hue. But owing to the relatively weak S cone influence, responses to the intermediate as well as 0 and 180 deg. hues were at least as strong as those to 90 or 270 deg.

To more directly assess the predictions from mixing of COFDs to generate color opponent signals, we also generated DKL hue tuning curves for the pixels at the intersections corresponding to the regions where neurons would be expected to be shifted toward each of the red (L+ ∩ S+), yellow (L+ ∩ S-), blue (M+ ∩ S+), and green (M+ ∩ S-) color appearance mechanisms (Fig. 6g and Extended Data Fig. 4f). As expected from their L+ contributions, the “red” (e,g. L+ ∩ S+) and “yellow” pixels were strongly biased to preferences toward 0 deg. and away from 180 deg. hues. Similarly, the pixels corresponding to “colors” with M+ contributions (“blue” and “green”) had DKL hue preferences that were strongly biased toward 180 deg. and away from 0 deg. Importantly, it can be seen that the differential mixing of S+ and S- for each of the combinations generates differential shifts toward or away from the S+ (90 deg.) and S- (270 deg.) DKL hues. Specifically, the “red” pixels have hue preferences that are shifted toward 90 deg. (away from 270 deg.) relative to the “yellow” pixels, and the “blue” are similarly shifted relative to the “green” pixels. Again, despite the clear differences between the intersections with S+ versus S- COFD contributions, the shifts along the 90-270 axis are much smaller than the shifts along the 0-180 deg axis, once again reflecting the weak overall contributions of the S cone inputs. This imbalance can also be visualized by comparing to an idealized schematic that assumes balanced interactions across the COFD intersections (Fig. 6h) to combine cone-opponent signals. Fig. 6h also serves to illustrate the hypothesized production of hue pinwheels at some locations where the physical positioning of COFDs generates interactions across orthogonal S+/S- and L+M−/M+L- axes (Figs. 4h,i and 6a-c). In such cases, the conceptualized schematic and the actual physical mapping on the brain have similar configurations.

### COFDs and CO histology

Previous studies have suggested that cytochrome oxidase blobs in layer 2/3 are specialized for processing of color information^23^ (but see^10,46^), receive direct S+ input from the LGN^9,47^, and could have neurons with a different distribution of preferred colors than interblobs^29^. We were therefore interested in whether there might be a relationship between COFDs and CO staining, either for all COFDs together (data from all 5 animals) or for ON or OFF domains of particular cones (animal A7 excluded). Alignment of COFD maps to postmortem CO histology shows that they have similar periodicity and are often overlapping, but there are also clearly regions that do not overlap (Fig. 7e,f,h and Extended Data Fig. 7). To quantify these relationships, the distributions of CO intensities were compared between pixels located within versus outside all COFDs. Complete results from all comparisons for all imaging regions are shown in Extended Data Fig. 7a. In general, regardless of the cone type (L, M, S) or phase (ON or OFF), pixels in the COFDs have higher CO intensities and pixels in non-COFD regions have lower CO intensities compared to the overall distribution of CO intensities across V1 (Fig. 7g). The biases of these COFD regions toward high CO intensities were all statistically significant for every animal (Extended Data Fig. 7a, 4 animals for comparisons between phase-identified COFDs with non-COFDs, 5 animals for comparisons between phase-unassigned COFDs with non-COFDs). These statistical comparisons within animals are based on treatment of each COFD region as an independent sample (Wilcoxon signed rank test), and *P* values less than 0.05 are considered significant. Across animals, the bias toward high CO intensities is statistically significant (*n* = 5 animals, *P* = 0.03125, right-tailed Sign test) only for the comparisons based on phase-unassigned COFDs (5 animals), as maximal statistical significance for 4 animals is *P* = 0.0625. The achromatic ON and OFF domains were also significantly biased toward blob regions for all 3 animals in which achromatic phase maps were generated.

**Fig. 7.**
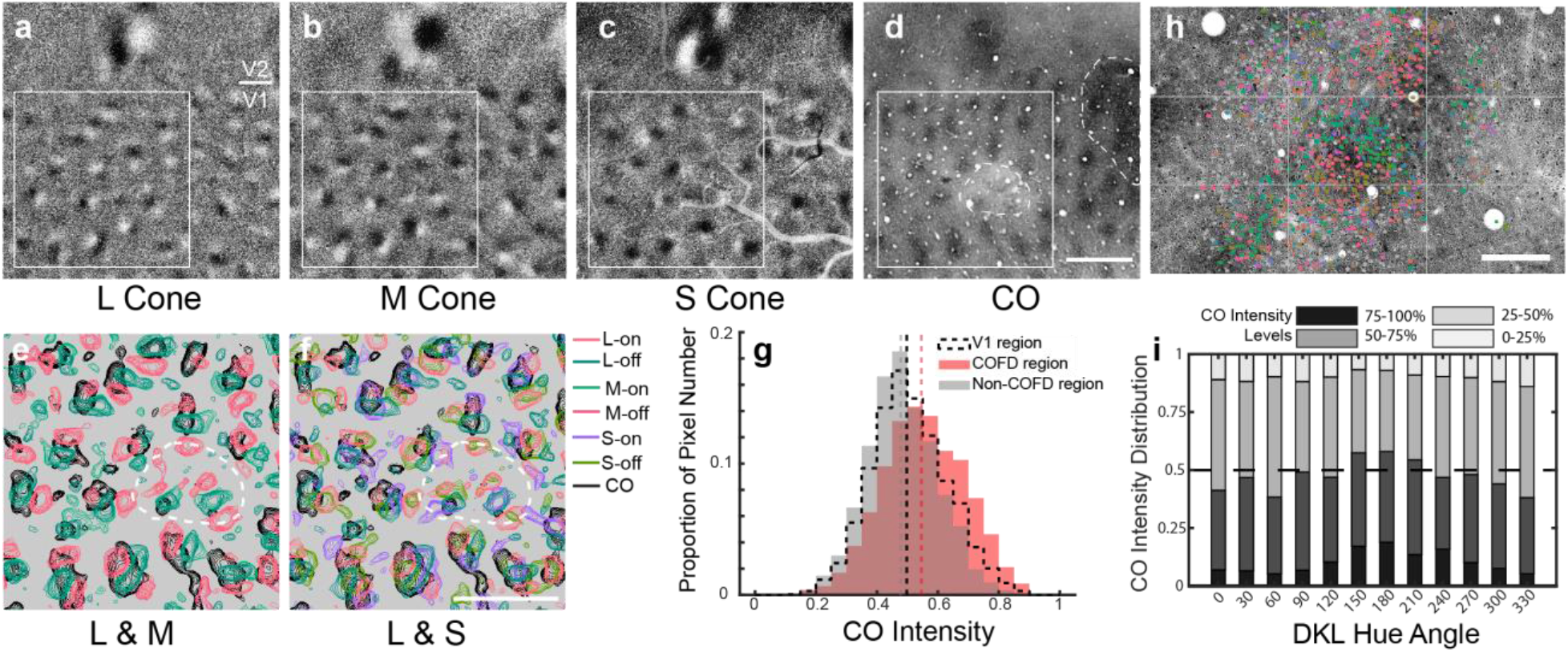
Spatial relationship between COFDs and CO blobs. **a-c**, L- (**a**), M- (**b**), and S-cone (**c**) COFD maps of V1 and V2 from animal A5. **d**, CO image after alignment. It is the same region shown in **a-c**. Scale bar in **d**: 1 mm; applies to **a-d**. **e-f**, Zoom-in view of the COFD and CO contours from the region within the white square shown in **a-d**. More cases are shown in Extended Data Fig. 7a. Scale bar in **f**: 1 mm; applies to **e** and **f**. **g**, Histogram of CO intensity distributions in COFD, non-COFD and whole V1 regions. Dashed lines are median of the distribution in each region. The CO intensity distribution in COFD regions is significantly higher than that in non-COFD regions (*n* = 96 COFD regions, *P* = 0.0062, Wilcoxon signed rank test). Note that dotted white outline in **d-f** indicates region restricted from quantitative analysis due to damage by an electrode penetration (subsequent to ISI data collection) and loss of CO staining. **h**, DKL hue preference map overlays on top of cytochrome oxidase (CO) histology. Scale bar in **h**: 200 μm. Note that h is the same region shown in Fig. 3f-i and Fig. 5f-i. Grids were added as landmarks for comparison. **i**, CO intensity distribution of DKL hue preference neurons. Each column is the CO intensity distribution (grouped as four levels) of pixels corresponding to the neurons (based on the alignment shown in **h**) prefer the hue direction shown on *x* axis. Results are from 5 regions (10 planes, see Extended Data Fig. 5); **h** is one example.

Because a previous study has shown that S+(L+M)-LGN afferents preferentially terminate in the CO blobs^9^, we were particularly interested in whether S+ domains might be more strongly biased toward high CO intensity regions than other COFDs. Comparisons between cone types and phases within animals revealed consistent significant differences only for the S+ domains, which were significantly more biased toward higher CO intensities than S- for all four animals (Wilcoxon rank sum test, *P* values less than 0.05 are considered significant, and the same test method and criteria were applied on the following comparisons). S+ was also significantly higher than both L+ and M- in 3 of 4 animals. Other comparisons were significant for only 1 or 2 out of the 4 animals. It is noteworthy that the densest S+ LGN terminals and CO staining are found deep within layer 2/3 (layer 3B)^9,47^, so there might be a closer correspondence between S+ responses and CO deeper in cortex.

We also quantified the relationships between CO staining and the preferred directions in DKL color space of neurons imaged with 2PCI (Fig. 7i). The highest CO staining intensities were observed at the locations of neurons preferring color directions around 180 deg. (greenish) while the lowest CO intensities were observed at the locations of neurons preferring color directions around 0 deg. (“pink”). These observations are consistent with the relationships previously reported between CO staining and color preferences of neurons tested using CIE colors^29^; neurons preferring green were observed to be at locations of higher CO staining intensity than those preferring red.

Previous studies have used ISI and colored visual stimuli to reveal patchy activation patterns (“color domains”) within V1^26,28^. We have generated ISI maps using the same methods applied in those studies (regions preferentially responsive to red/green or blue/yellow isoluminant stimuli) and directly compared those maps to COFDs imaged in the same animals. We find that the color domains and COFDs are closely related (Extended Data Fig. 7). This is expected, because both red/green and blue-yellow isoluminant modulation should differentially activate selected regions within the COFDs. The partial but not exclusive correspondence between COFDs and CO histology is also consistent with previous observations of the relationships between color domains and CO staining^28^.

## Discussion

In summary, we have demonstrated that: 1) L-, M-, S-COFDs and achromatic ON/OFF domains are a prominent feature of the functional organization of primate V1 that can be readily revealed with ISI imaging; 2) 2PCI-assayed neurons within ON/OFF domains of L, M, S and achromatic have the same ON/OFF sign of that domain type; 3) both qualitative and quantitative assessment of the spatial intersections of COFDs shows that they follow the mixing rules predicted from the Three-stage model; 4) DKL hue phase maps show that the hue preferences at intersections between COFDs correspond to hues predicted from functional mixing and generate hue pinwheel or linear structures; 5) 2PCI of DKL hue preferences at the intersections of COFDs shows that functional mixing occurs at the level of individual neurons; and 6) COFDs are biased toward regions of high CO intensity. We conclude that the neural substrates that underlie the mixing required for a transition from cone-opponent (stage 2) mechanisms to color opponent (stage 3) mechanisms are highly organized and implemented by specific connections between neurons in the geniculate input layers (4C, 4A, 3B) and the neurons that we have assayed in more superficial V1.

The most prominent feature of Stage 3 in the Three-stage model is the specific mixing between cone-opponent mechanisms that are predicted to give rise to each of the four color-opponent mechanisms (Fig. 1). This mixing is expected to generate a greater range of hue preferences than is present in the cone-opponent geniculate input population, largely owing to the interactions between the L/M and S/(L+M) mechanisms that do not occur at earlier stages. That this mixing begins in V1 is expected from previous studies evaluating color preferences of V1 neurons^10^. The “unique colors” (red, green, blue and yellow) are not necessarily expected to be preferred by larger numbers of neurons than other hues, but instead reflect perceptual transitions within the color-opponent mechanisms, where, for example, “red” appears neither “bluish” or “yellowish”^1^. (Due to the opponent nature of color appearance, red cannot appear “greenish”^19^.) Consistent with these expectations, our observations and analyses indicate that the mixing of L/M and S/(L+M) opponent signals within COFDs gives rise to shifts in preferred hues that are in the directions expected to generate color-opponent mechanisms. But the contribution of the S/(L+M) mechanism (S-cone contributions) is likely much less than expected to fully account for color appearance. Careful calculations based on psychophysical measures assign just a 2.55-fold stronger influence to the L/M mechanism^1^, while the distribution of color preferences we have measured from neurons in upper layer 2/3 of V1 appear to reflect a much stronger relative influence of the L/M mechanism (Fig. 5n). It is likely that these relative contributions are re-weighted in higher visual areas, as suggested by a recent 2PCI study comparing responses to CIE hues in cortical areas V1, V2, and V4^48^. Importantly, our observations using DKL hues show that the biases of V1 neurons toward end-spectral CIE colors observed previously^29,48^ do not result solely from the stronger L and M cone activation generated by the red and blue CIE hues^29,49^; S-cone contributions are weak even in response to DKL stimuli with matched L, M, and S-cone increment magnitudes. Neurons deeper in layers 2/3 and 4A (inaccessible to 2PCI) that receive direct S/(L+M) signals^9^ and can project directly to V2 might also have more dominant S-cone influences that could contribute to stronger S cone influences in higher visual areas.

We were surprised to find that ISI mapping of the locations preferring DKL hues modulated in different directions (hue direction maps) revealed “pinwheel” structures (Fig. 6) reminiscent of previously described orientation^38,40–43^ and direction maps^39,44,45^. Hue direction pinwheels were surrounded by regions of iso-hue direction, again reminiscent of iso-orientation and iso-direction domains. Hue direction pinwheels were often found at the intersections of L/M opponent and S- cone COFDs and in the most striking cases aligned with the intersections of orthogonal L+M−/M+L- and S+/S- COFD pairs (e.g. Figs. 6 a-c and schematized in Fig. 6h). Additional pinwheels were also located outside of the COFDs. We speculate that hue direction pinwheels emerge through developmental mechanisms that favor the grouping of neurons with similar functional preferences interacting with the requirement for retinotopic mapping (e.g. through wire length optimization^50^) and may share common principles with those underlying emergence of orientation and direction maps.

Electrode penetrations extending across cortical layers have also found locations containing neurons with Blue-Yellow versus Red-Green opponency, leading to the suggestion that there are separate blobs for each type of opponency^24^. The COFDs that we observe, as well as their relationships to CO staining, argue against this interpretation and provide an explanation for why different electrode penetrations might sometimes be biased to sample neurons with different color opponency. There may in fact be color columns that extend across cortical layers, but they likely correspond to the more closely spaced COFDs with different cone inputs rather than to separate blobs (see further below). Future experiments using ISI imaging of COFDs to guide subsequent electrode penetrations (e.g., Extended Data Fig. 9) should more clearly reveal the extent to which color selectivities and/or cone input signs and sensitivity extend across or vary between layers.

We have observed a functional organization (COFDs) in the most superficial layers of V1 that involves systematic mixing of L/M opponent with S+ and S- signals. The fact that L/M opponent LGN inputs terminate in a separate layer (layer 4Cβ) from S+ (layer 3B) and S- inputs (layer 4A) indicates that mixing of these cone-opponent systems can only occur through neural substrates that span across these layers and also suggests that there are likely to be some laminar differences in cone responses and color tuning. The most prominent substrate for potential mixing is the axons of layer 4Cβ spiny stellate neurons that extend to and densely arborize in layers 4A and 3B, likely mixing L/M opponent with S cone signals^51,52^. Yet another synaptic step is required to transmit information in layer 4A/3B to the neurons that we imaged in layer 2/3A^53^. An additional possibility that should not be overlooked is a specialized population of layer 6 pyramidal neurons (Type IβA) that have both dendritic and axonal processes that arborize selectively in both layers 4Cβ and 4A/3B^54–56^. These neurons could directly integrate L/M opponent with S+ or S- LGN inputs and provide their mixed signals to neurons in layers 4Cβ and 4A/3B. Such a mechanism could account for Blue-Yellow opponent neurons that have been reported in layer 4C^24^. While our observations clearly point to these circuits as the neural substrates that mediate mixing of cone-opponent mechanisms to generate the rudiments of color opponency, further experiments will be required to more precisely identify the micro-circuitry involved.

The V1 functional organization and micro-organization related to cone-opponency and color preferences that we have reported here build on a long history of prior studies using varied technologies and sometimes leading to apparently contradictory results. However, in the light of the higher resolution and denser sampling afforded by 2PCI technology, sources of the apparent discrepancies are now much clearer. Before the development of ISI, studies used CO blobs as an anatomical landmark for linking the functional properties of sparsely sampled neurons to possible functional architecture. Notably, Livingstone and Hubel first reported a prevalence of neurons in and around CO blobs that were color selective but not orientation selective, suggesting that color tuning would likely have some sort of functional organization. (But see^57^.) This was later reinforced by ISI studies demonstrating patchy cortical activation with iso-luminant, full-field colored stimuli ^26,28^. While the overall organization of color patches was similar to CO blobs with substantial overlap, there was clearly not a one-to-one correspondence. Those observations clarified that when single unit recordings are related to blobs, blobs are an indirect and imperfect proxy for color functional architecture. Other studies combining single unit recordings with postmortem CO histology provided the first evidence for a columnar organization for color preference and suggested that separate blobs might process “red-green” (L/M) versus “blue-yellow” (S/L+M) opponency ^24,25^. But the first ISI maps of V1 locations preferring different colors suggested color transitions on a finer spatial scale than would be expected from separate L/M and S/(L+M) blobs ^27^. This was confirmed when 2PCI was used to definitively identify functional micro-architecture for color preferences at the level of single neurons ^29,30^. Neurons preferring similar colors are grouped together within a functional color microarchitecture and neurons preferring different colors are spaced more closely than would be compatible with their separation by blobs.

Both Garg et al.^29^ and Chatterjee et al.^30^ also present results indicating that color mapping is dependent on mapping for spatial features, as first suggested by Livingstone and Hubel^23^. Chatterjee et al.^30^ show that color selective neurons responding to full-field stimuli are strongly biased toward blobs, while those in the inter-blobs are relatively suppressed by full field stimuli and instead respond well to drifting gratings that contain oriented edges. Garg et al.^29^ report that non-orientation-selective, color-preferring neurons are strongly biased toward blobs, but orientation-selective color-preferring neurons are not.

Using CIE colors, Garg et al.^29^ also noted that neurons preferring green were biased much more strongly toward high CO regions than those preferring red or blue. Consistent with those observations, we find higher CO staining at locations with neurons preferring DKL hue directions around 180 deg (“greenish”) than 0 deg. (“pink”). Here we also report that all of the regions within COFDs (generated with full-field stimuli) are significantly biased toward high CO activity (“blobs”) and that S-ON regions are significantly more biased than S-OFF COFDs. Altogether, these results point to an organization in which the blobs are a reliable marker of locations receiving direct S-ON input from the LGN^9,47^ and are part of a somewhat larger, overlapping system of COFDs. The COFDs mediate an orderly mixing of the S-ON input to blobs with L+/M- and M+/L-input from layer 4Cβ and S-OFF input from layer 4A, that follows the stage 3 mixing rules to generate a broader range of color preferences arranged in hue pinwheels.

## Methods

### Animals and animal care

All procedures involving live animals were conducted in accordance with the guidelines of the NIH and were approved by the Institutional Animal Care and Use Committee (IACUC) at The Salk Institute for Biological Studies. We recorded from seven adult macaque monkeys (*M. fascicularis*) under anesthesia in this study. They are referred as A1 to A7. Details and allocation of these animals can be found in **Extended Data Table 1**. In brief, two-photon calcium imaging (2PCI) data were collected from A1, A3 and A4. Electrophysiology data were collected from A2 and A5. Intrinsic signal optical imaging (ISI) and histological data were collected from A1, A2, A5, A6 and A7.

### Surgery

Surgical procedures for AAV injections (recovery surgery), physiological recordings (non-recovery) and euthanasia in this study were the same as described previously^29^.

Briefly, for animals used for 2PCI (A1, A3 and A4), two sequential surgeries were performed. First a recovery surgery was conducted for injection of AAVs to express the genetically-encoded calcium sensor, GCaMP6f (see below). Then after 10-12 days recovery to allow for GCaMP expression, a second non-recovery surgery was performed and an imaging chamber was implanted for ISI and 2PCI. For other animals used in this study, only the non-recovery surgery was performed.

Before all surgeries, animals were anesthetized with ketamine (10 mg/kg) and pretreated with atropine sulfate (0.02 mg/kg) intramuscularly. During surgeries, animal were under general anesthesia (1-2.5% isoflurane in O2) and sterile conditions. Animals were intubated (tracheostomy was performed during non-recovery surgery) and mechanically ventilated. EKG, SpO2, end tidal CO2, and body temperature were all monitored throughout the procedure and during all recordings. Lactated Ringer’s with 5% dextrose, antibiotics (Cefazolin, 25 mg/kg, i.v.), and dexamethasone (1-2 mg/kg, i.v.) were supplied.

During the first surgery, following a craniotomy over V1, a 1:1 mixture of AAV5-TRE3-GCaMP6f and AAV5-thy1s-tTa^58^ (Salk Viral Core GT3, titer: 8.04E+12 GC/ml and 2.13E+12 GC/ml) was pressure injected via fine glass micropipettes at 1-3 depths in each of 6-12 locations in V1 (300-600 nl per injection, 300-700 μm below cortical surface). Injections were spaced about 1mm apart to yield nearly continuous labeling. Following the injections, a thin piece of silicone artificial dura was placed between the cortical surface and the natural dura and these were glued in place with Vetbond (3M, Maplewood, MN). The bone was replaced and cemented in place. Animals were recovered. Buprenorphine was administered for three days (slow release, 0.12 mg/kg, s.c.). Antibiotics (Cefazolin, 25 mg/kg, i.m., one week) and dexamethasone (1-2 mg/kg, i.m., three days) were administered daily to prevent infection and reduce brain swelling.

In the non-recovery surgery, a head-post was implanted to stabilize the animal. A large imaging chamber (20 mm inner diameter) was cemented onto the skull over V1 and V2. A coverslip insert was screwed into the chamber to reduce brain movement. Wires were implanted through small hole in the skull and connected to an amplifier to monitor EEG. Anesthesia was gradually transitioned from isoflurane to sufentanil citrate (2.0-8.0 μg/kg/hr, i.v.), and vecuronium bromide (0.05-0.2 mg/kg/hr i.v.) was administered to paralyze the animal before recording and EEG was subsequently monitored to continuously along with other vital signs to assess anesthetic depth. Anesthesia dosage was adjusted at a level that insured slow wave EEG activity. Eyes were dilated with 1% atropine sulfate and 0.5% proparacaine hydrochloride ophthalmic solution and fitted with contact lenses of appropriate curvatures to focus on a screen 57 cm from the eyes. First, we used ISI to obtain functional maps (e.g., ocular dominance map, orientation map, cone-opponent phase map, etc.) to target the regions of interest (e.g., cone-opponent functional domains) during 2PCI (on animals A1, A3 and A4, described below) or extracellular electrophysiology recording (on animals A2 and A5, described below). The animals were humanely euthanized by anesthetic overdose followed by perfusion at the end of the procedure (see “Histology and Image Alignment”).

### Intrinsic Signal Optical Imaging (ISI)

Surface blood vessel images were acquired under 540 nm illumination. Images of reflectance change (hemodynamic signals corresponding to cortical activity) were acquired under 630 nm illumination. Imaging was performed using a GigE camera (Photonfocus, Switzerland) and custom Matlab software.

In this study, we conducted two ISI paradigms known as standard episodic stimulation combined with blockwise data acquisition and continuous-periodic stimulation combined with continuous data acquisition^36^. (see “Visual Stimulus Presentation”). For creating ocular dominance, orientation and isoluminant color maps, we utilized the first ISI paradigm. Images were acquired during the pre-stimulus and stimulus periods (6 seconds, see below) at 10 Hz. For cone-opponent and DKL hue-phase maps, we utilized the second ISI paradigm, images were acquired at 10 Hz during stimulus presentation, which consists of a 2-second pre-stimulus period and a 10-20 minute stimulus period.

### Two-photon Microscopy

Microscopy for two-photon calcium imaging (2PCI) was performed using a customized setup incorporating a Sutter movable objective microscope (Sutter Instruments, Novato, CA) with a resonant scanner (Cambridge Instruments, Bedford, MA) and data acquisition controlled by a customized version of Scanbox (Neurolabware, Los Angeles, CA). GCaMP6f was excited by a Ti:sapphire laser (Chameleon Ultra II, Coherent, Santa Clara, CA) at 920 nm. Of three animals used for 2PCI, two (A3 and A4) were single-plane continuously scanned at 15.49 Hz (unidirectional scanning); one (A1) was double-plane continuously scanned with an electrically tunable lens (Optotune, Switzerland) at 30.98 Hz (bidirectional scanning). An area of about 1168 μm × 568 μm (A3 and A4) or 1100 μm × 725 μm (A1) was imaged with a 16x, 0.8 NA objective lens (Nikon Corporation, Tokyo, Japan). The back aperture of the objective was not overfilled due to the technical limitations of the microscope.

### Extracellular Electrophysiology

Extracellular electrophysiological data were collected using Neuropixels Phase 3 probes with 384 recording channels^59^. Briefly, signals were sampled at 30 kHz and filtered separately to an action potential band (0.3–10 kHz) and a local field potential band (LFP, 0.5–1000 Hz), then streamed to a computer under the control of SpikeGLX (http://billkarsh.github.io/SpikeGLX) software. The probe was held and lowered into the cortex using an oil hydraulic manipulator (Narishige International USA, Inc.), which was attached to the imaging chamber in order to ensure perpendicular penetration to the brain surface through a small hole in the coverslip. The locations of the electrode penetration were within cone-opponent functional domains (COFDs) revealed by ISI, and the probe was slowly lowered 2.5-3.1 mm below brain surface. The brain surface at the location of the penetration was covered with 2% agarose and then with silicone elastomer (Sylgard 184, Dow Corning, USA) to prevent drying. Data were recorded at least 30 min later after the electrode was stable and visual fields were subsequently mapped.

### Visual Stimulus Presentation

Visual stimuli were generated with the Psychophysics Toolbox Version 3 (http://psychtoolbox.org/) for MATLAB (Mathworks Inc., MA) on a CRT monitor (CPD-G520, Sony Corporation, Japan) with a refresh rate of 100 Hz. For all experiments, the monitor was positioned 57 cm in front of the animal’s eyes. Prior to each experiment, the luminance of each of the red, green and blue phosphors was linearized. Then a white point and mean luminance were chosen (*x* = 0.30, *y* = 0.31, *Y* = 71.5 cd/m^2^ for A1; *x* = 0.32, *y* = 0.34, *Y* = 86.3 cd/m^2^ for A2) as the background for spectra measurement of each phosphor with a spectroradiometer (PR-701, Photo Research, Syracuse, NY). Then we utilized the spectra of each phosphor to calculate gun gains for cone-isolating stimuli (including Hartley flash gratings and temporal-modulating gratings, see below) and DKL hue stimuli (see below, including drifting gratings and temporally-modulated gratings, see below), and we used the same white point and mean luminance as background, inter-stimulus interval and blank condition during cone-isolating stimuli, DKL hue stimuli and full contrast achromatic grating stimuli (including drifting gratings and temporal-modulated gratings, see below) presentation. Another white point (*x* = 0.33, *y* = 0.33, *Y* = 10 ± 0.1 cd/m^2^ for other animals) was chosen as the background and blank condition for physical equal luminance CIE (Commission internationale de l’éclairage) hue stimuli as previously published^29^ (also referred to as CIE hues below).

For ISI experiments, both episodic and continuous-periodic stimuli were utilized. Episodic full-field achromatic square-wave drifting gratings (8 directions, 1.5 cycles/degree, 6.67 cycles/sec, duty cycle: 0.2) were presented binocularly to locate orientation columns and monocularly to obtain an ocular dominance (OD) map. Four CIE hues (red and green, blue and yellow) were chosen to construct Red/Green and Blue/Yellow isoluminant square-wave drifting gratings (4 directions, 0.2 cycles/degree, 2 cycles/sec, duty cycle: 0.5) to obtain Red/Green and Blue/Yellow isoluminant color maps. In order to increase the signal-to-noise ratio, each stimulus condition was presented in a randomized order with 20 to 30 repeats. For each trial, the stimulus was displayed for 4 seconds with a 2-second pre-stimulus period and 8-second inter-stimulus interval. Continuous-periodic full-field cone-isolating (**Fig. 3a**), achromatic, and DKL hue temporally modulated gratings (**Fig. 5a**) were only modulated at a low temporal frequency (0.1, 0.08 or 0.017 Hz) to obtain cone-opponent, achromatic ON/OFF and DKL hue-phase maps. The temporal profile of these cone-isolating and achromatic stimuli was sine-wave (**Fig. 3a**); the temporal profile of DKL hue stimulus was square-wave with 0.5 temporal duty cycle (**Fig. 5a**).

For 2PCI, only episodic stimuli (drifting and Hartley flashed sine-wave gratings) were used and all stimuli were presented monocularly. First, the dominant eye was selected based on the ISI ocular dominance map of the imaging area; the non-dominant eye was occluded. All stimuli were given through the dominant eye. Next, the receptive field of the imaging area was mapped by showing a series of drifting gratings and the stimulus size (1.3-1.5 degree in diameter) was chosen to be as small as possible while still covering receptive fields of all neurons within the imaging region.

Drifting gratings (achromatic, DKL hue and CIE hue) were presented for 3-4 seconds with a 3- to 4-second inter-stimulus interval. Achromatic drifting gratings were presented as sine-wave gratings at 8 directions (0 to 315 degrees in increments of 45 degrees), 5 or 6 spatial frequencies (0.2, 0.4, 0.8, 1.6, 3.2, 6.4 cycles/degree), and a single temporal frequency of 5 cycles/sec. Each condition was repeated 5 times in random order. DKL and CIE hue drifting gratings were presented as square-wave gratings with 0.5 spatial duty cycle, 8 directions (same as achromatic gratings), 4 spatial frequencies (0.2, 0.4, 0.8, 1.6 cycles/degree), and a single temporal frequency of 5 cycles/sec. Each condition was repeated 3 times in random order.

The Hartley stimulus (flashed sine-wave gratings) set^31^ consisted of 120 blank stimuli and 5640 unique gratings with different combinations of orientation, spatial frequency (range: 0.2-6.0 cycles/degree), and spatial phase (range: 0, 90, 180, 270 degree) (**Fig. 2a**). The stimulus was presented for 60 trials (60 seconds per trial), and between each trial there was a 2 second inter-stimulus interval. Within each trial, stimulus conditions were randomly selected from the stimulus set with replacement and were flashed continuously at 4 stimuli per second. A TTL pulse was generated at each stimulus transition and time-stamped with the corresponding microscope scanning frame and line number. The time stamps were used to synchronize stimulus and imaging frames for reverse correlation analysis (see “Spike-Triggered Average and Cone Weight Calculation”). The Hartley stimulus set was generated with L, M, S cone-isolating colors and 98%-99% contrast achromatic color.

For extracellular electrophysiology recording, a uniform patch was temporally modulated along two axes in the DKL isoluminant plane at 0.33 Hz with 0.5 temporal duty cycle (see “DKL Hues” below). The patch was presented monocularly to measure the ON/OFF responses to verify the ON/OFF phases (see “Intrinsic Signal Optical Imaging Data Analysis”) of the cone-opponent and DKL hue-phase maps.

### Cone-isolating Calculation

We followed a previously described method of calculating cone-isolating directions ^32^. Briefly, we calculated a 3 × 3 transformation matrix (*M*) by taking the dot product of the monitor spectral power distribution (*r*(*λ*), *g*(*λ*), *b*(*λ*). *λ* was sampled in the range [380, 780]nm every 2 nm) of the 10° cone fundamentals (*l*(*λ*), *m*(*λ*), *s*(*λ*)) published by Stockman and Sharpe^60^.

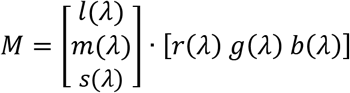

Next, we utilized the following equation to calculate the R, G, B gun gains for each L, M, S cone-isolating direction. Then, the gun gains for each direction were normalized to [−1 to 1].

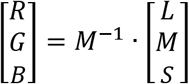

In all experiments, we matched the cone contrast of the L and M cone-isolating stimuli and ran the S cone-isolating stimuli at the maximum achievable contrast. The monitor was re-calibrated prior to each experiment. Consequently, the exact contrasts of the cone-isolating stimuli changed slightly between experiments (L: 18.0%-18.5%; M: 18.0%-18.5%; S: 89.9%-90.2%).

### DKL Hue

The calculation of the DKL color space was followed as previously described by Brainard^61^. The background (center of DKL isoluminant plane) was the same background we used for measuring phosphor spectra (see “Visual Stimulus Presentation”). The same cone fundamentals used for cone-isolating stimuli were used for DKL hue calculation. Twelve hues were selected from the DKL isoluminant plane with equal increments from 0 to 330 degree (**Fig. 5a and Extended Data Fig. 8b**). For A1, the total cone-increment (pooled differential L-, M- and S-cone excitation between DKL hue and background) of each DKL hue was matched (**Figs. 5f-i, 7h, Extended Data Figs. 4a, 4d, 5, 8b, and Extended Data Table 2**); for A2, each DKL hue had the maximum achievable cone contrasts by our display (**Figs. 5b-e**, **6b, 6c, Extended Data Figs. 4a**, **8b**, and **Extended Data Table 2**). Each cycle of spatial (for 2PCI) or temporal (for ISI) DKL hue square-wave gratings consists of a DKL hue and the background color with a 0.5 duty cycle. Except for this monopolar temporal modulation of DKL hues (modulation between DKL hues and the background) used for ISI, two bipolar temporal modulation of DKL hues (modulation between two DKL hues) were also applied for electrophysiology only. One of the bipolar modulations used two hues along L-M axis (0 and 180 degree, representing L+M- and M+L-respectively) in DKL isoluminant plane. Another utilized two hues along S-(L+M) axis, which are 90 and 270 degrees in DKL isolumiant plane, representing S+(L+M)- and (L+M)+S-, respectively.

### CIE Hues

The CIE hue stimuli were identical to those described in a previous publication^29^; more details of the stimuli can be found in the supplementary materials of that paper. Briefly, twelve hues were selected from the HSL plane (S = 100%, L = 50%) with equal increment from 0 to 330 degree. Then the CIE-1931 *xyY* coordinates (**Extended Data Fig. 8a and Extended Data Table 2**) of each hue were measured with a spectroradiometer, and we adjusted the R, G, B value of each hue to match their luminance (*Y*) to 10 ± 0.1 cd/m^2^. Because of this adjustment and the gamut of the CRT monitor, the twelve hues were not in the same plane of the HSL color space, but they had the same luminance (*Y*). The CIE *xy* coordinates of each hue remained the same between different experiments. For A3 and A4, we presented the CIE hues. One cycle of the CIE hue spatial square-wave gratings consists of a CIE hue and the background color (*x* = 0.33, *y* = 0.33, *Y* = 10 ± 0.1 cd/m^2^, see “Visual Stimulus Presentation”) with 0.5 duty cycle.

### Histology and Image Alignment

Histology was performed as previously described^29^. Animals were euthanized by a lethal dose of pentobarbital sodium (Euthasol, i.v.) and transcardial perfusion was performed using 0.9% saline, followed by 4% paraformaldehyde (PFA), 10% sucrose with PFA and 20% sucrose with PFA, and the brain was extracted. The recording region was then blocked, flattened and sunk in 30% sucrose and sectioned (50 μm) tangentially using a freezing microtome. Tissue was stained for cytochrome oxidase (CO) and sections were imaged. Both CO and 2PCI images were aligned manually with the ISI blood vessel map in Adobe Photoshop. Regions from ISI images corresponding to 2PCI images were interpolated in Adobe Photoshop using ‘Bicubic Automatic’ method.

### Intrinsic Signal Optical Imaging Data Analysis

Data was analyzed by customized software written in MATLAB. For episodic stimulation paradigm, percent change (*ΔR*/*R*) of each pixel for each stimulus condition was first calculated using the following formula:

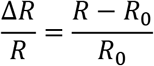

*R* is the average response of a pixel during stimulus period, and *R*0 is the average response of a pixel during pre-stimulus period. The *ΔR*/*R* image of each condition was then filtered using a fourth-order Butterworth bandpass filter (high cutoff frequency: 7.8-11.2 cycle/mm, low cutoff frequency: 0.8-2.1 cycle/mm). Next, we computed a t-map, as previously described^39^, and based on the *ΔR*/*R* images of two conditions to reveal the OD columns and orientation columns. The calculation of orientation polar map (from t-maps, **Extended Data Fig. 5**), DKL hue direction (from DKL hue-phase maps, **Fig. 6b,c and Extended Data Fig. 4d**), and DKL hue polar map (hue direction map scaled by vector magnitude pixel-by-pixel, **Fig. 5g and Extended Data Fig. 5**) was the same as described previously^38,39^. For continuous-periodic stimulation paradigm, the phase map computation have been described^36^. Specifically, the *ΔR*/*R* image of each camera frame during stimulus period was first calculated to reduce noise, then the phase of each pixel was extracted from the *ΔR*/*R* image sequence to plot phase map. Finally, the phase map was clipped to ±1.5 SD of the image, and then filtered using contrast-limited adaptive histogram equalization (CLAHE) method provided by MATLAB. Hemodynamic delay in phase map was not compensated as previously described^36^; instead, we took the advantage of the results from 2PCI or electrophysiology recording to identify the sign of ON/OFF phases in the ISI phase maps (**Figs. 3, 4, 5h-i, 7a-c**, **and Extended Data Figs. 1**, **2b, 3a-b, 9b**).

### Image Processing for COFD Contours, Histograms and ISI DKL hue tuning curves

To further investigate the spatial relationship between different COFDs and between COFDs and DKL hue domains, COFD contours, bivariate histograms and the DKL hue tuning curve of each COFD were plotted. To plot COFD contour maps, COFD phase maps were first filtered with a 2-dimensional Gaussian kernel (σ = [5, 5]), then contours were extracted at different pixel-value levels from filtered images to generate the COFD contour maps (only top and bottom 15% contours were shown, **Figs. 4d-h, 6a-b, 7e-f**, **and Extended Data Figs. 2a**, **3b-c, 7a-b, 9b**).

To plot bivariate histograms, COFD masks were generated by selecting one level of contours if the size of the contour is close to the domain size in phase maps. We manually removed some regions (e.g., blood vessel, noise) to make sure the masks are specific to regions of interest. A blood vessel mask and area mask were also added to exclude pixels within blood vessel regions or other areas outside V1 (e.g., V2). Next, COFD phase maps and DKL hue phase maps were filtered with a 2-dimensional Gaussian kernel (σ = [5, 5]) and normalized to [−1 to 1]. Then pixels on COFD maps were selected based on the combined L, M, S cone COFD masks to plot bivariate histograms to show spatial relationship between different COFD maps (**Fig. 4j-l and Extended Data Fig. 2a**). Similar histograms were also plotted to show the spatial relationship between COFD maps and Achromatic ON/OFF map (**Extended Data Fig. 3d-f**). Pixels on COFD maps and DKL hue-phase maps were selected based on each L, M, S cone COFD mask to plot bivariate histograms to show spatial relationship between COFDs and DKL hue domains (**Fig. 6d** and **Extended Data Fig. 4b**). All bivariate histograms have 50 × 50 bins; bin counts of each histogram were normalized to [0 to 1]. Specifically, for each COFD-DKL hue bivariate histogram, the center of top 10% bright pixel was calculated in up-quadrant and bottom-quadrant independently. Then, the angle of the line connecting centers in up- and bottom-quadrants was calculated to show the L-, M- and S-COFDs have different and systematic spatial relationship with DKL hue domains (**Fig. 6e** and **Extended Data Fig. 4c)**. Curves were fitted using MATLAB fit function with smoothing spline model.

To plot the DKL hue tuning curve of each COFD, DKL hue phase maps were filtered with a 2-dimensional Gaussian kernel (σ = [5, 5]) and normalized to [0 to 1]. The mean pixel value of each DKL hue-phase map was calculated based on each L, M, S cone COFD mask (**Fig. 6f** and **Extended Data Fig. 4e**) or based on the intersection regions of COFD masks (**Fig. 6g** and **Extended Data Fig. 4f**). Curves were fitted by Generalized von Mises function (*GvM*) as described by the following equation^62,63^:

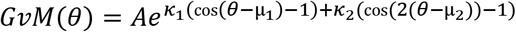

Where *θ* is the DKL hue direction in the range of [0 2π); μ_1_ ∈ [0 2π) and μ_2_ ∈ [0 π) are preferred directions; κ_1_, κ_2_ > 0 are the measure of concentration about μ_1_ and μ_2_.

To plot the spatial relationship between CO intensity and COFD maps, blood vessel holes in CO images were first filled using ‘Content-Aware’ method in Adobe Photoshop. Then CO images were filtered with a 2-dimensional Gaussian kernel (σ = [5, 5]) and normalized to [0 to 1] with higher pixel value means higher CO intensity. Pixels within COFD regions (combined L-, M-, S-cone COFD masks) and non-COFD regions were selected (pixels within blood vessels were excluded) to plot the histogram (**Fig. 7g and Extended Data Fig. 7a**). If there was bleach of CO intensity in CO image due to laser scanning or electrode penetration (e.g., **Fig. 7d**), pixels within that region were excluded for quantitative analysis. After aligned DKL hue preference map with CO image (**Fig. 7h**), pixels corresponding to neurons preferring the same hue were grouped, and the CO intensity distribution of that group was plotted as 4 levels (0-25%, 25%-50%, 50%-75% and 75%-100%) in one column shown in **Fig. 7i**. Pixels within each COFD and achromatic ON and OFF mask, and within whole V1 region were selected and grouped into 4 levels (0-25%, 25%-50%, 50%-75% and 75%-100%) based on CO intensity levels, see **Extended Data Fig. 7a**. For animal A7, only histograms of COFD regions versus non-COFD regions were plotted because we did not identify the ON/OFF phase of the COFDs from that animal (**Extended Data Fig. 7a**).

### Two-photon Calcium Imaging Data Analysis

Regions of interest (neuron cell bodies) were manually segmented using Adobe Photoshop. Raw fluorescence signals were extracted and the spike rate was estimated by Scanbox (https://scanbox.org/) as previously described^64^. For two-plane bidirectional scanning, raw fluorescence signals were interpolated before spike estimation. Thus, interpolated signals and spikes had 30.98 Hz sampling rate for both planes. Once signals and spikes were extracted, they were analyzed using customized software written in MATLAB and The Julia Language (https://julialang.org/). The change in fluorescence *(ΔF/F*) was computed by the following equation, where *F* is the average fluorescence during stimulus presentation and *F*_0_ is the average fluorescence during the pre-stimulus period:

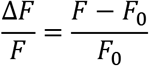

Responses were averaged over 3-5 stimulus presentations, and the stimulus condition (e.g, direction, spatial frequency and hue) of the maximal response was identified. Neurons were considered visually responsive if they had a *P* < 0.05 to either two-way ANOVA across all stimulus conditions (including blank stimuli), or two-tailed Welch’s *t*-test between the strongest response and response to blank stimulus.

To determine whether a neuron was significantly direction and/or orientation tuned, we chose all direction conditions at a neuron’s preferred spatial frequency. Then we applied circular variance method^65,66^ to average the responses of 8 direction to get 1 direction vector. Each condition was repeated 5 times in the experiment, so there were 5 direction vectors for each neuron. Similarly, 5 orientation vectors were obtained by averaging two directions of motion at each orientation. Then we projected the 5 vectors onto their final mean vector, thus the 2-dimensional distribution of 5 vectors was transformed to a 1-dimensional distribution. Note that each of those 5 vectors depended on how the conditions combined as one repeat, therefore the condition combination affected the distribution of 5 vectors. To reduce the distribution bias introduced by low sampling, we randomly shuffled direction or orientation conditions for 100 times and calculated the vectors to produce 500 data points demonstrating the distribution of the direction or orientation response magnitude of a neuron. We did the same procedures for the blank condition and calculated the distribution of the blank response magnitude of the same neuron. Then we analyzed the two distributions using ROC analysis^67^ and calculated the probability of the area under a ROC curve (AUC, used the trapezoidal rule) as a parameter to indicate the significance of direction or orientation selectivity. If AUC > 0.7, we considered the neuron to be direction or orientation selective. The preferred direction or orientation is the angle of final mean vector from the circular variance calculation (**Extended Data Fig. 5**).

To determine whether a neuron was significantly hue tuned, we first tested whether the neuron was direction and orientation tuned as described above, but chose all directions using the spatial frequency and hue to which the neuron was maximally responsive. If a neuron was direction or orientation tuned, all hue conditions were chosen under the neuron’s preferred spatial frequency and direction or orientation. Otherwise, hue conditions were chosen under the neuron’s preferred spatial frequency for circular variance and ROC analysis, similar to determination of direction and orientation selectivity. The AUC was calculated based on the ROC curve of distributions of blank and hue conditions. If AUC > 0.8, we considered the neuron to be hue selective, and the preferred hue was the hue to which the neuron maximally responded (**Fig. 5f** and **Extended Data Fig. 5**).

To show the relationship between neurons’ DKL hue preference and COFD organization, neurons within COFD regions were plotted as dots labeled with their preferred hues with normalized L-, M-, and S-cone COFD magnitude as coordinates (**Fig. 5j-l**). First, 2PCI images were aligned with the ISI blood vessel map, and regions of ISI images corresponding to 2PCI images were interpolated (see “Histology and Image Alignment”). Then, interpolated ISI images were filtered with a 2-dimensional Gaussian kernel (σ = [20, 20]) and normalized to [−1 to 1]. Next, pixels on L-, M-, and S-cone COFD maps were selected based on the neuronal segmentation mask to calculate the normalized COFD magnitude for pixels corresponding to each neuron. Neurons within COFD regions from 5 imaging regions (10 planes) of A1 were selected to plot neuronal COFD magnitude distribution with their preferred DKL hues (**Fig. 5j-l**).

### Spike-triggered Average and Cone Weight Calculation

First, we used the time stamps to synchronize the stimulus sequence and estimated spike trains (**Fig. 2**). Then, the linear kernel of receptive field was estimated by spike-triggered average (STA) using following formula modified from previous publication^68^:

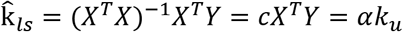

Where 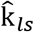 is the estimated kernel using least-square, *Y* is a column vector of neural responses to each unique stimulus condition been presented and *X* is the stimulus matrix with each row is linearized unique stimulus. All 4 Hartley stimulus sets (L, M, and S cone-isolating, and achromatic) share the same Hartley basis function, which is radially symmetric, therefore, (*X*^*T*^*X*)^−1^ can be simplified as constant *c*, which is the same for all Hartley stimulus set. Note that this constant *c* is canceled when calculate magnitude change and does not affect the normalized cone weights calculation neither (see below). The estimated kernel 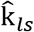 is a 2D image, which is also a vector in multidimensional space with each pixel as a dimension. The vector 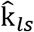 can be represented as its magnitude (*α*) multiples with its unit vector (*k*_*u*_). STA was calculated using 33 ms (bidirectional scanning) or 66 ms (unidirectional scanning) time bins of estimated spikes within a time window (−400 ms to 66 ms). In this STA movie, we defined a neuron’s best kernel estimation (*k*_*ubest*_) as the time when the magnitude reached its peak (*α*_*max*_), and this time was defined as best time delay. Typically, this occurred 198-300 ms prior to a spike, which is very similar to the time window reported in mouse visual cortex^69^. To check the significance of *k*_*ubest*_, we defined the kernel estimated in the time window (0 ms to 66 ms) after each spike as ‘blank’ kernel (*k*_*ublank*_) with its magnitude (*α*_*ublank*_), then z-scored *k*_*ubest*_ was computed based on *k*_*ublank*_. The *k*_*ubest*_ was considered significant if it passed all three criteria: 1), its *α*_*max*_ must within 198 ms to 300 ms time window prior to a spike; 2), its magnitude change (Δα/α, defined below) must bigger than 0.25 threshold; 3), the variance in z-scored *k*_*ubest*_ was 3.5x greater than the variance of z-scored *k*_*ublank*_, or the absolute mean of z-scored *k*_*ubest*_ was 3.5x greater the absolute mean of z-scored *k*_*ublank*_.

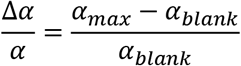

A neuron had 4 best kernels calculated from L, M, S cone-isolating and achromatic Hartley stimulus sets, respectively; if any of 4 kernels passed those three criteria, this neuron was considered as having a kernel that can be estimated by the STA of that stimulus set. A neuron could have one or more significant kernels from different Hartley stimulus sets.

If a neuron has a significant kernel to a Hartley stimulus set, we obtained the sign (positive or negative) of the maximal absolute value in its best kernel, and used the sign to plot the ON (positive) and OFF (negative) maps (**Figs. 2, 3, and Extended Data Fig. 1**).

To calculate the normalized cone weights in **Fig. 5m** and **Extended Data Fig. 6a**, only neurons with a significant kernel to the L, M, or S cone-isolating Hartley set were included. For each neuron in this group, we first compared the *α*_*max*_ from three cone types, and we defined the dominant cone type as the one with maximal *α*_*max*_. Then, we used the best time delay of the dominant cone type to slice out three images (*k*_*u*_) from L, M and S cone STA movies. Next, we made a mask by thresholding the top 5% of absolute pixel values in the image of the dominant cone type, and used the same mask on all three images to select pixels to calculate their mean value. The mean values from L, M, and S cone images are the raw cone weights (*w*_*l*_, *w*_*m*_, *w*_*s*_, respectively). The normalized L cone weight (*W*_*l*_) was calculated using the following equation, and similarly for the normalized M and S cone weight (*W*_*m*_ and *W*_*s*_).

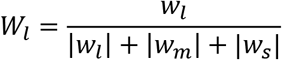

In the L-M cone weight plots provided (**Fig. 5m** and **Extended Data Fig. 6a**), the S cone weight is implicit in the distance from the diagonal lines. We chose neurons with a significant kernel to S cone-isolating Hartley set, and plotted the histograms based on the ON/OFF signs to S cone stimuli and their preferred hues (**Fig. 5n-p** and **Extended Data Fig. 6b-d**).

### Extracellular Electrophysiology Data Analysis

Data were analyzed using customized software written in The Julia Language. The LFP gamma band power has been reported to be closely related to hemodynamic signals recorded by ISI^70^. Therefore, we utilized the LFP gamma band power to identify the ON/OFF phases of the COFD phase map. The LPF band (0.5-1000 Hz, see “Extracellular Electrophysiology”) was bandpass filtered (1-100 Hz) and 60 Hz line noise was removed. Then, LPF power spectrums were estimated by Multi-Taper methods from 0-300 ms following and −300-0 ms before stimulus onset which correspond to the response (*P*_*r*_) and baseline (*P*_*b*_) epochs. Stimulus responses were defined as relative change of the power spectrum (*P*_*rc*_) within these epochs using following equation.

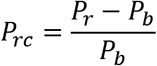

The gamma band power (30-100 Hz) of the recording channels within superficial layer (0-300 μm) were chosen to identify ON/OFF phases of the COFD within which electrode penetrations were made (**Extended Data Fig. 9**).

### Statistics

In all statistical tests, *P* < 0.05 was considered significant. The visual responsiveness of neurons was tested using two-way ANOVA and two-tailed Welch’s *t*-test. The distributions of neuronal DKL hue preferences were compared using Fisher’s exact test.

To statically compare the CO intensity distribution within COFD regions versus non-COFD regions, COFD and non-COFD regions were selected using masks described above. For within animal comparisons, each COFD region was treated as an independent sample, which was represented as the median of the CO intensity within that COFD region. For each animal, the CO intensity median values (*x*) within COFD regions were compared with the median (*y*) within non-COFD regions to test whether (*x*-*y*) has zero median against the alternative hypothesis that the median is not zero using Wilcoxon signed rank test. For comparison across animals, we applied Sign test (right-tailed) to see whether the difference between the median of CO intensity within all COFDs and non-COFD is significant across animals. The same Wilcoxon signed rank test was also performed on achromatic ON/OFF regions versus non-COFD region. For animals in which we identified the ON/OFF sign of ISI phase map, we compared the CO intensity distribution in ON regions versus OFF regions within and between each cone type in each animal. The L-, M-, and S-COFD masks were the same as described above. We performed Wilcoxon rank sum test to check whether two distributions have equal medians.

## Data and Code Availability

All data and code necessary to support the paper’s conclusions are present in the main text, supplemental information, or available from the corresponding author upon reasonable request.

## Acknowledgments

We thank Professor Tetsuo Yamamori’s laboratory for kindly providing pAAV-TRE3-GCaMP6f and pAAV-thy1s-tTa to make AAVs used in the study. We thank the Callaway lab for support, particularly Drs. B. J. Hansen, E. J. Kim and E. Richler. We thank Drs. D. L. Ringach, A. Cheng, R. Liu, and K. J. Nielsen for their assistance and suggestions with our two-photon microscope. We thank Dr. T. P. Franken for his contribution to the ISI acquisition software. We thank Dr. I. Nauhaus for his contribution to the visual stimulus code. This work was supported by NIH grants EY022577 (E.M.C), NS105129 (E.M.C.), MH063912 (E.M.C.), EY028084 (A.K.G), the Gatsby Charitable Trust (E.M.C.), and the Pioneer Fund (P.L.).

## Author contributions

P.L., A.K.G. and E.M.C. designed the experiments. P.L. designed and built the ISI setup. P.L. designed the imaging chamber. A.K.G. built the electrophysiological recoding setup. P.L. and E.M.C. performed recovery surgeries and virus injections with M.S.R. and A.K.G.’s assistance. P.L. contributed to the visual stimulus code. P.L. performed non-recovery surgeries and set up the ISI, 2PCI and electrophysiological recoding with M.S.R. and A.K.G.’s assistance. P.L., A.K.G., M.S.R. and L.A.Z. acquired the ISI, 2PCI and electrophysiological data. P.L. and M.S.R. performed the CO histology and scanned the slides. P.L. and A.K.G. contributed to the ISI data analysis code. L.A.Z., P.L. and A.K.G. contributed to the 2PCI data analysis code. P.L. analyzed the ISI, 2PCI and histology data and made the figures. L.A.Z. contributed to the electrophysiology data analysis code, analyzed the data and made the figure. E.M.C. and P.L. wrote the manuscript with contributions from all other authors.

## Competing interests

The authors declare no competing interests.

---

**Extended Data Fig. 1.**
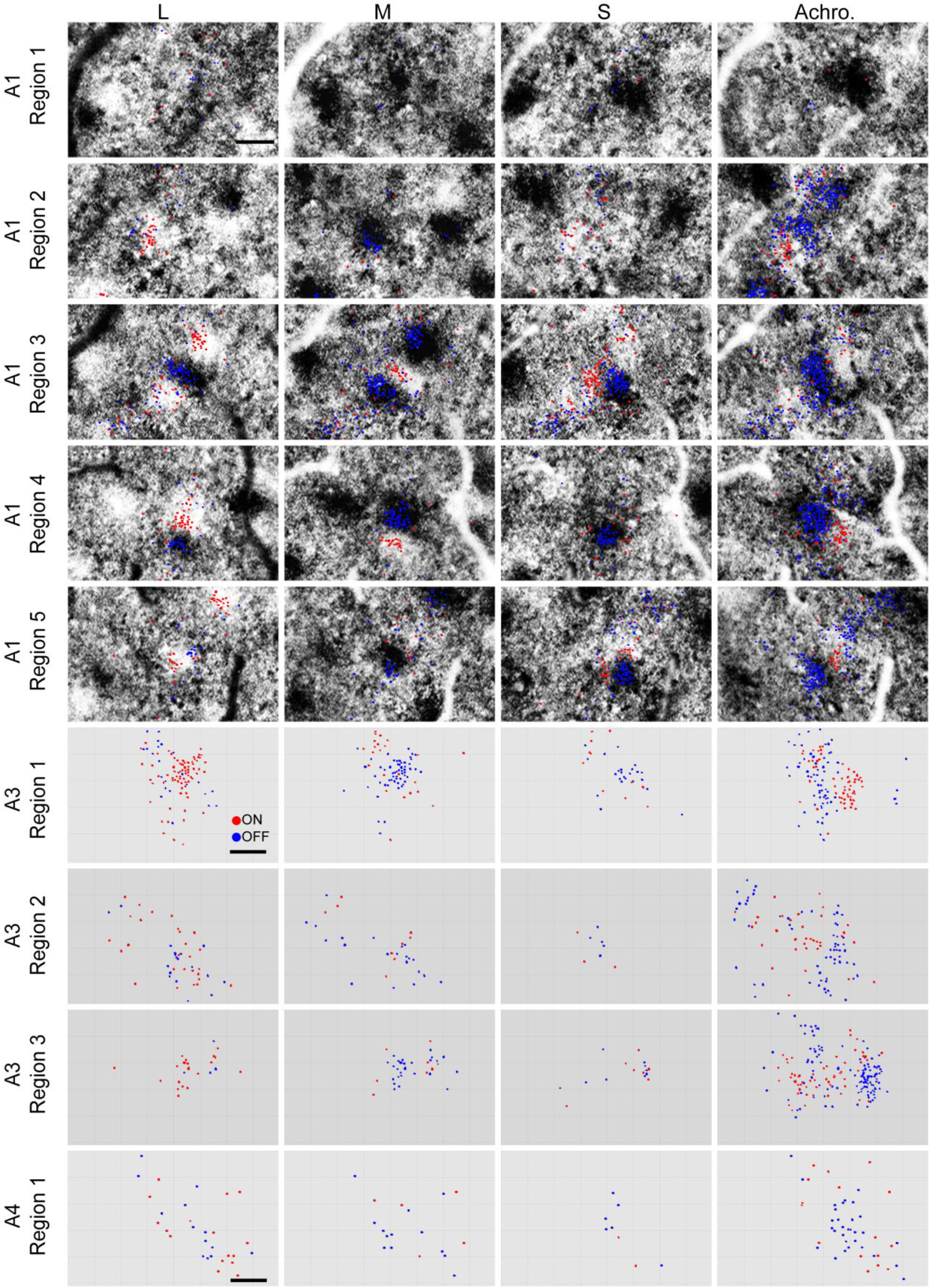
Additional cases of ON/OFF-dominant achromatic and cone-opponent receptive field clusters. Each row corresponds to one imaging region with neurons (dots) from two planes merged. Each column corresponds to responses to one type of cone-isolating (L, M, and S) or achromatic (Achro.) stimulus. Animals and imaging regions that were the source of the data are indicated to the left of each row. The first 5 rows are imaging regions from A1, with 2PCI neuron properties overlain on ISI COFD phase maps. “A1 Region3” is the same region shown in Figs. 2i, 3f-i, 5f-i, and 7h. “A1 Region4” is the same region shown in Fig. 2j. Red dots represent ON-dominant neurons; blue dots represent OFF-dominant neurons. Scale bars: 200 μm; the three scale bars each apply to all additional panels from same animal.

**Extended Data Fig. 2.**
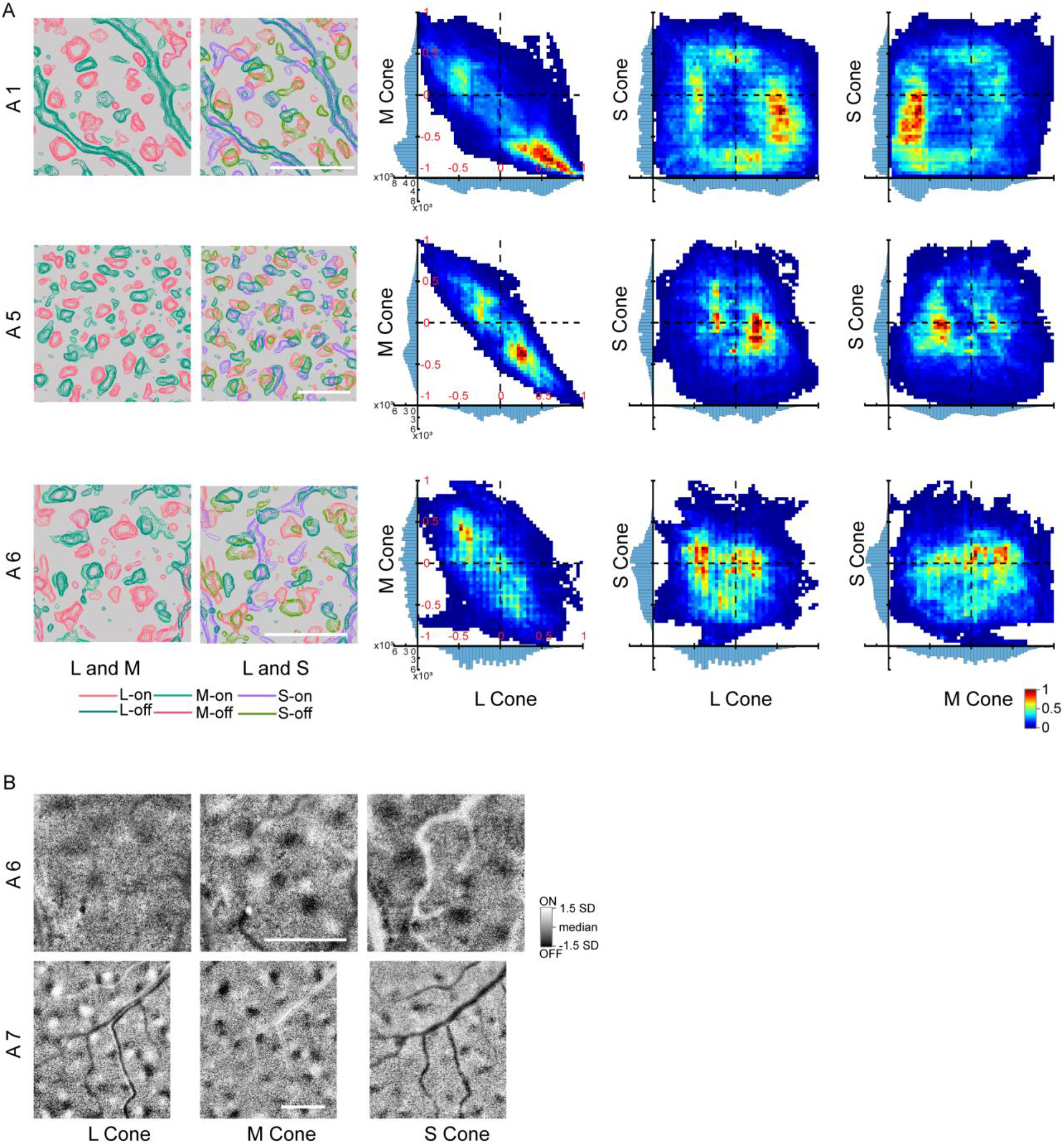
More cases of COFD maps and their spatial organization. **a**, Each row consists results from one animal. The COFD spatial organization and their relationship has been demonstrated in 4 ISI cases. One is shown in Fig. 4 (A2), others are shown here (A1, A5 and A6). First column is the overlay of L-cone and M-cone COFD contours. Second column is the overlay of L-cone and S-cone COFD contours. Scale bar for each case: 1 mm. Last three columns are the quantitatively analysis of the spatial relationship between L, M and S COFDs. **b**, Two more cases of COFD maps. For A7, we did not identify the ON/OFF phases of pixels in three maps. Scale bar for each case: 1 mm.

**Extended Data Fig. 3.**
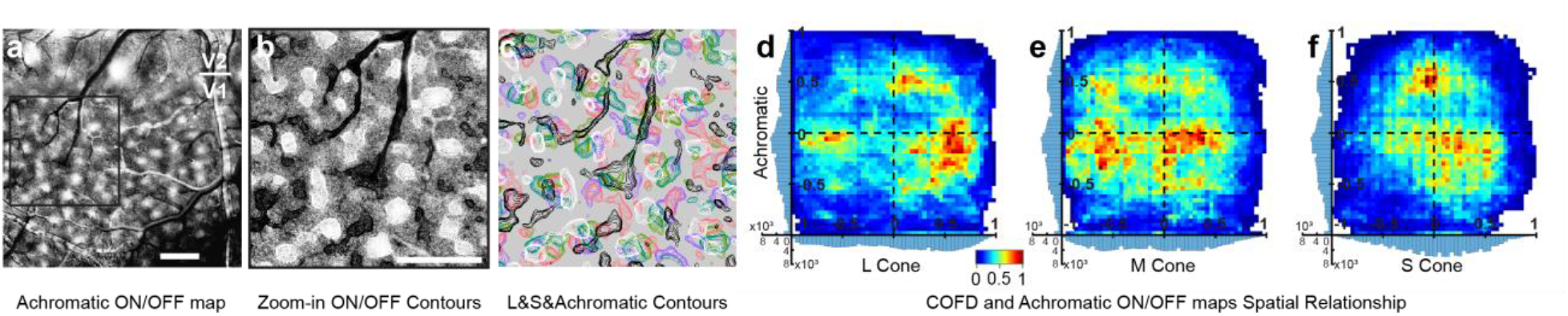
Achromatic ON/OFF map and its spatial relationships with COFDs. **a**, Achromatic ON/OFF map. Note that this is the same cortical region as shown in Fig. 4a-c. Scale bar: 1mm. **b**, Zoom-in view of the region selected in **a**, and overlaid with ON/OFF contours. Note that this is the same cortical region as shown in Fig. 4d-f. Scale bar: 1mm; applies to **b** and **c**. **c**, Overlays of the L- and S-COFD contours with Achromatic ON/OFF contours. **d-f**, 2D histograms of the spatial relationship between COFDs and Achromatic ON/OFF map. The 1D histograms on the *x* and *y* axes display the distribution of pixel values in each category.

**Extended Data Fig. 4.**
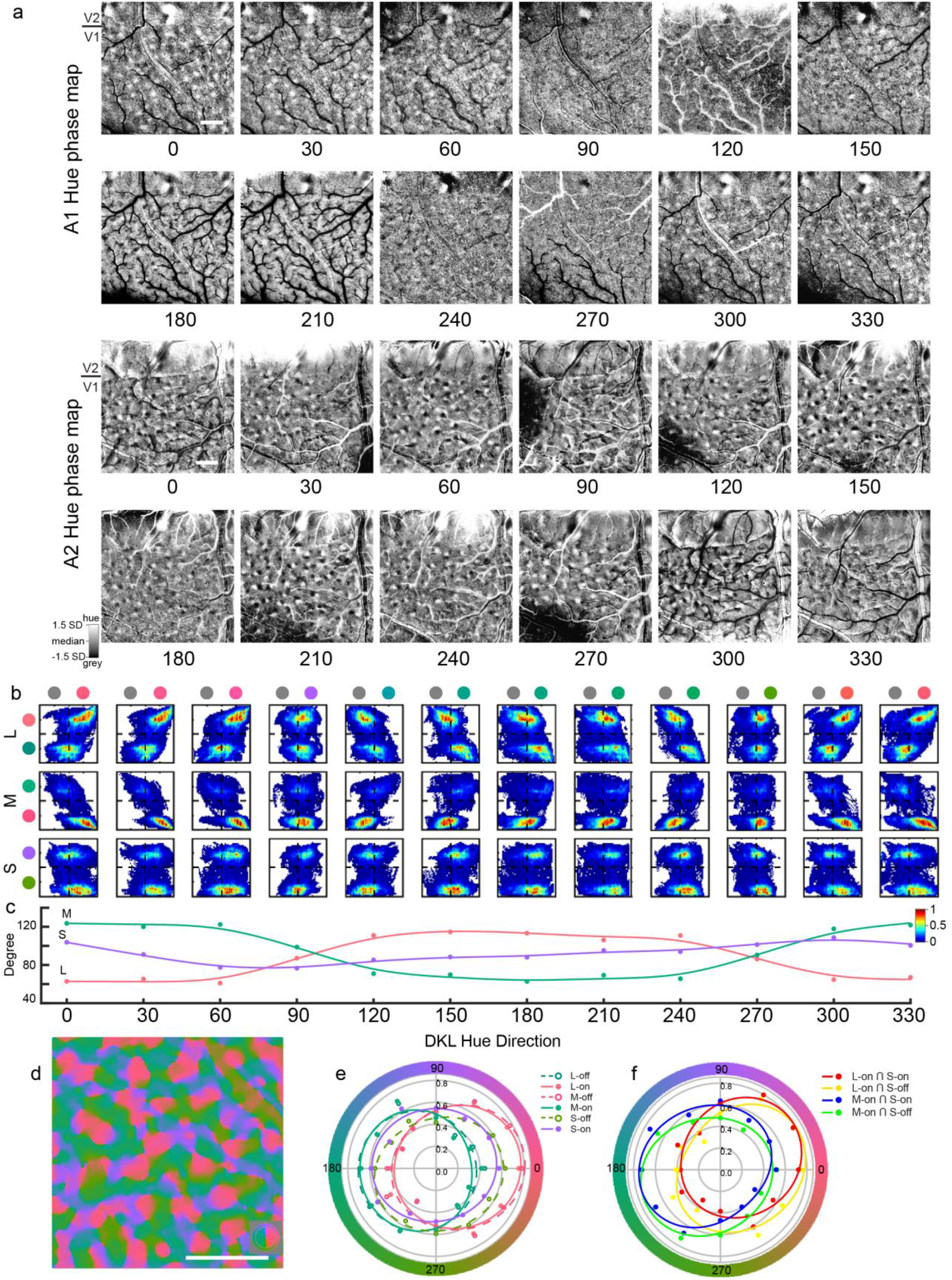
All DKL hue phase maps and their spatial relationship with L-, M-, and S-COFDs. **a**, All DKL hue phase maps from two animals (A1 and A2). The number under each image is the hue direction in DKL isoluminant plane. The DKL hues used for A1 were cone-increment-matched, in which S-cone modulation (90 and 270 degree) is weak, but for A2, cone contrast of each hue was maximized on the monitor; therefore, signals in A1 were weaker, especially along S-axis (90 and 270 degree). Scale bars in two cases are 1 mm. **b**, The bivariate histograms show systematic relationship between COFDs and DKL hue preference domains in animal A1. Same to Fig. 6d (A2), each row is from the same cone type, and each column is from the same DKL hue. On the left side of each cone type, there are two color disks, which represent the ON-phase (upper) and OFF-phase (lower), respectively. The color disks on top of each row represent the hue-phase and grey-phase, and the hue direction is labeled at bottom of **c**. **c**, The angle defined by the top 10% hot pixels in each bivariate histogram for each cone type. X-axis is the hue direction in DKL isoluminant plane, Y-axis is the angel calculate from each bivariate histogram (see “Image Processing for COFD Contours, Histograms and ISI DKL hue tuning curves”). **d**, DKL hue direction map of animal A1. The inserted color key is the same as the one shown in Fig 5f. The similar map from A2 is shown in Fig. 6b-c. Scale bar: 1mm. **e**, The hue tuning curves calculated from the mean of pixel values within each COFD region on hue preference maps in animal A1. **f**, The hue tuning curves calculated from the mean of pixel values within each COFD-overlapping region on hue preference maps in animal A1.

**Extended Data Fig. 5.**
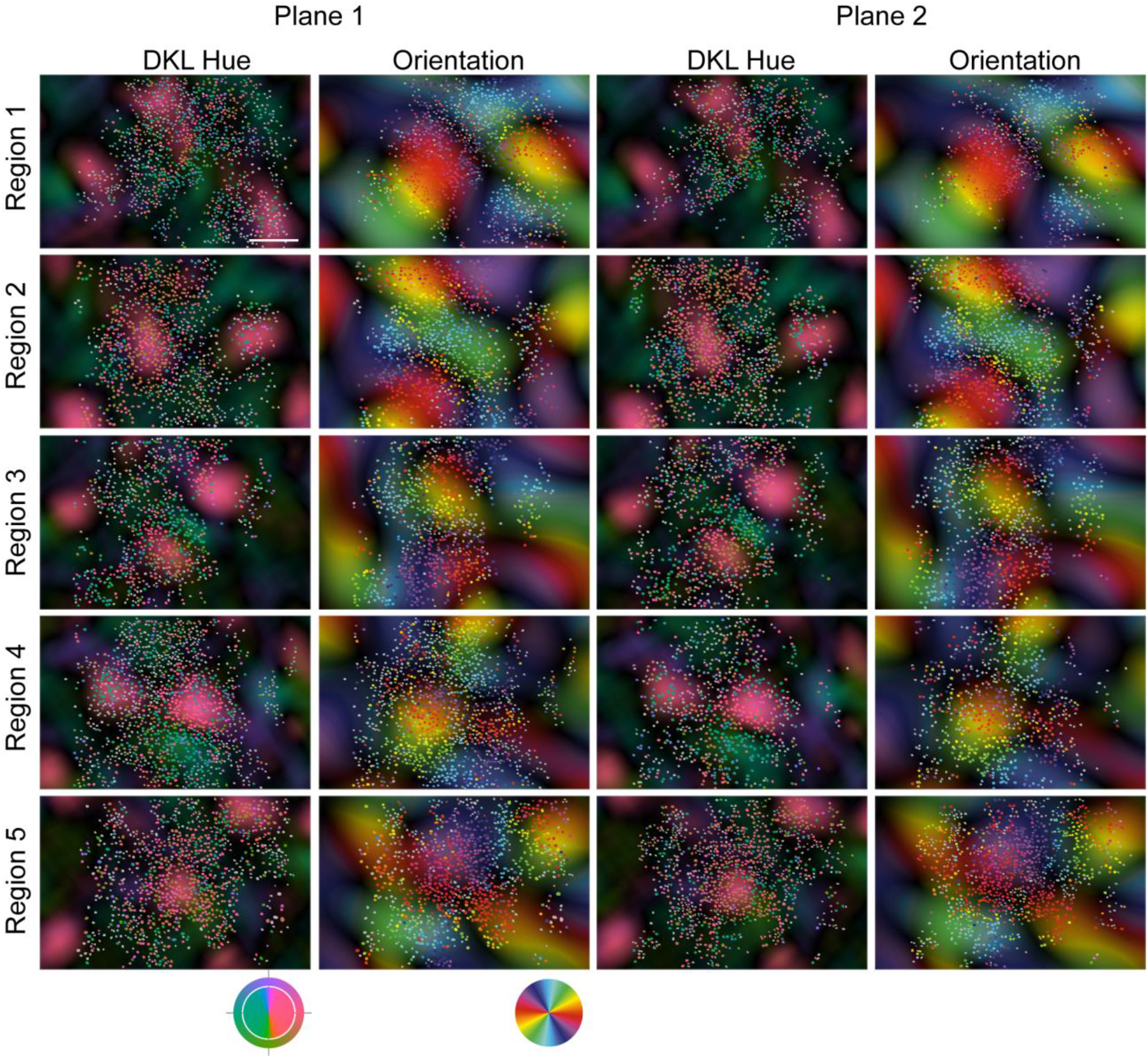
More cases of DKL hue preference maps and orientation maps. Each row is one imaging region with 2 planes. All 5 regions, as the same 5 regions shown in Fig. S1, are from A1. Each image has 2PCI image overlays on top of the polar map calculated from ISI images. Grey dot in 2PCI image indicates a neuron is not significantly tuned by DKL hue or orientation. Same to Fig. 3J, DKL hue preference maps of 2PCI and ISI are plotted using the DKL hue shown as the outer ring of the color key at bottom. The actual color used for stimulation is shown as the inner disk of the color key. Details of stimuli are shown in Methods. For orientation map of 2PCI and ISI, the orientation is indicated by the angle (0 to 180 degree) of color in the color key at bottom of the orientation map. We used the alignment of 2PCI orientation map and ISI orientation polar map as the secondary check about alignment between 2PCI and ISI maps based on surface blood vessel maps. Scale bar: 200 μm, applies to all images.

**Extended Data Fig. 6.**
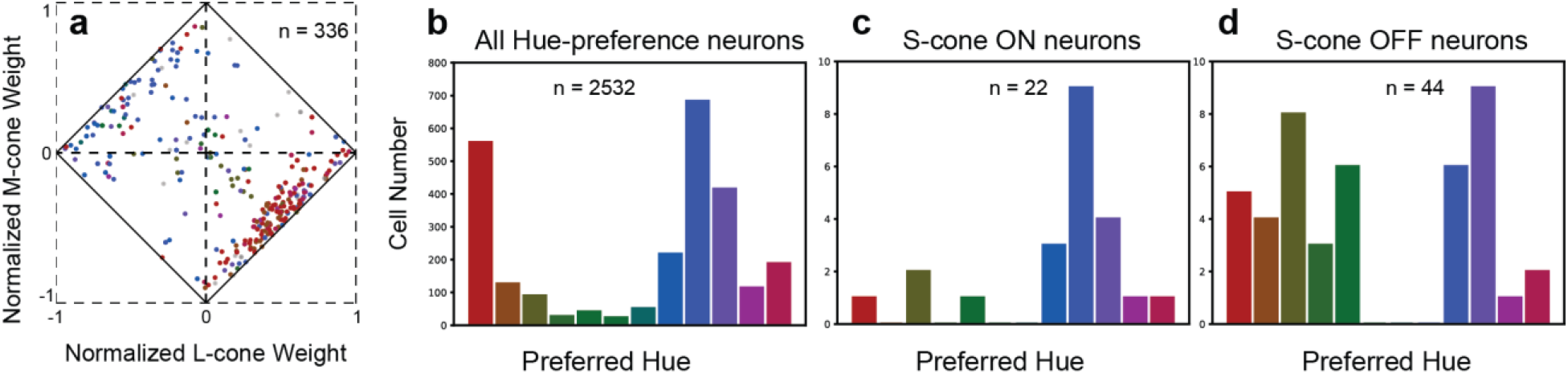
CIE hue preference and its relationship with Cone weights. **a**, Relationships between CIE hue preferences and cone weights calculated from STAs. 336 neurons with significant L, M, or S-cone STA kernels from 4 imaging regions (3 from A3 and 1 from A4, 6 planes total) are plotted. Each neuron in the plot is colored according to its preferred CIE hue. Neurons with significant STAs, but which are not hue selective, are plotted as grey. **b-d**, The distribution of preferred CIE hues of all hue-selective neurons (**b**), hue-selective neurons with significant S- ON STA (**c**), and hue-selective neurons with significant S-OFF STA (**d**). The preferred CIE hue is shown as the color of bar, and the number of neurons sampled is shown in each panel.

**Extended Data Fig. 7.**
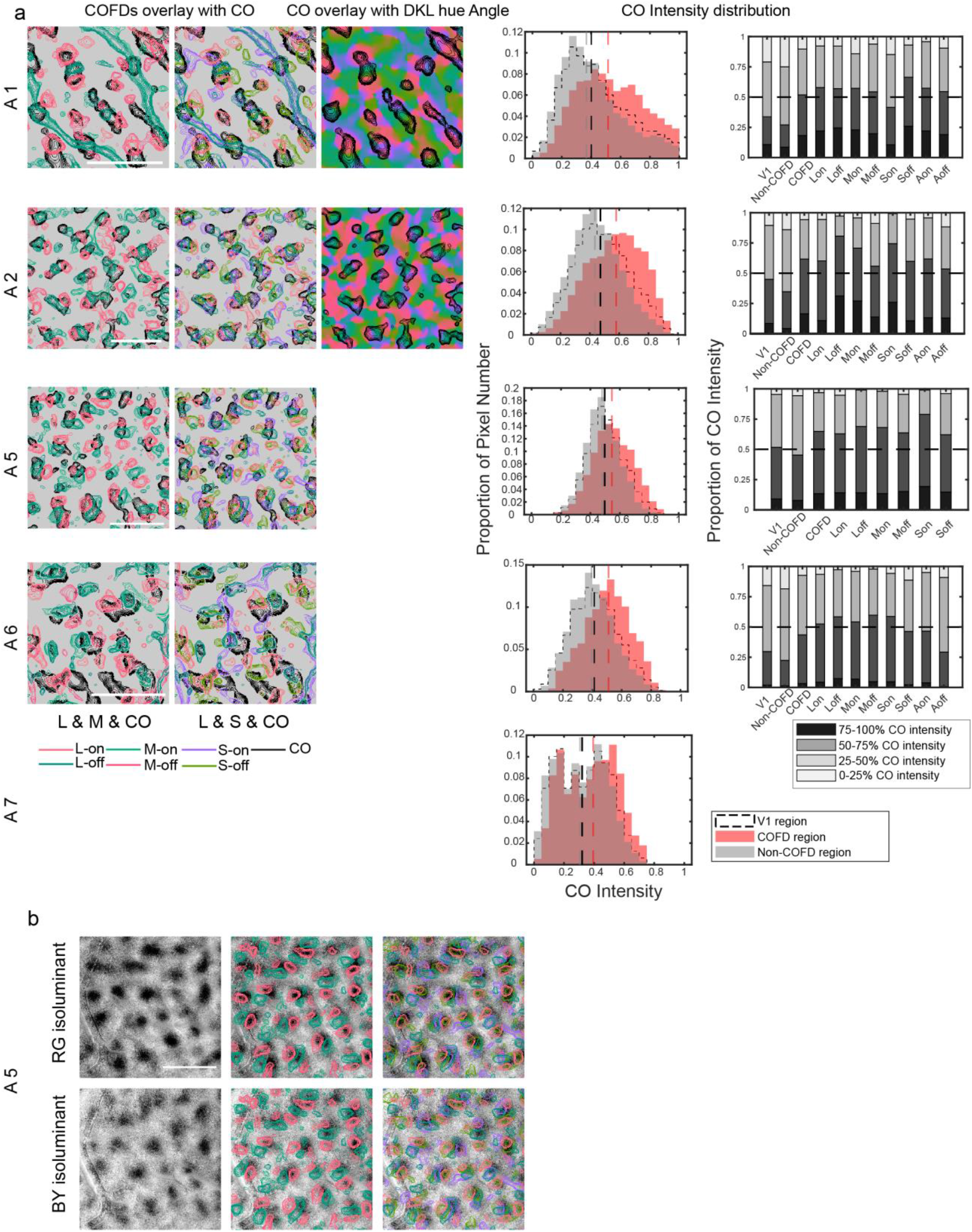
The spatial relationship of COFDs with CO blobs, DKL hue direction map, Red/Green and Blue/Yellow isoluminant color maps. **a**, Each row shows results from one animal. All 5 cases are shown in this Figure, with A5 also shown in Fig. 7. First column, L- and M-COFD contours overlay with CO blob contours. Second column, L- and S-COFD contours overlay with CO blob contours. Third column, which only consists of two cases (A1 and A2), is CO blob contours overlay on top of DKL hue direction map. Fourth column is the histograms of CO intensity distributions within COFD, outside COFD (Non-COFD) and whole V1 regions. Dash lines are median of the distribution in each region. In all cases, the CO intensity distribution is biased in COFD regions vs. Non-COFD regions, and the difference is statistically significant (Wilcoxon signed-rank test). Fifth column shows more comparison of the CO intensity distribution, including each COFD sub-region, achromatic ON (Aon) and OFF regions (Aoff). Note that because we did not identify the ON/OFF phase in A7, only comparison between COFD and Non-COFD was made. Scale bar for each case: 1 mm. **b**, COFD contours overlay on top of the Red/Green and Blue/Yellow isoluminant color maps. Note that these maps were from the same region of A5 in **a** and Fig. 7e,f. Scale bar for each case: 1 mm.

**Extended Data Fig. 8.**
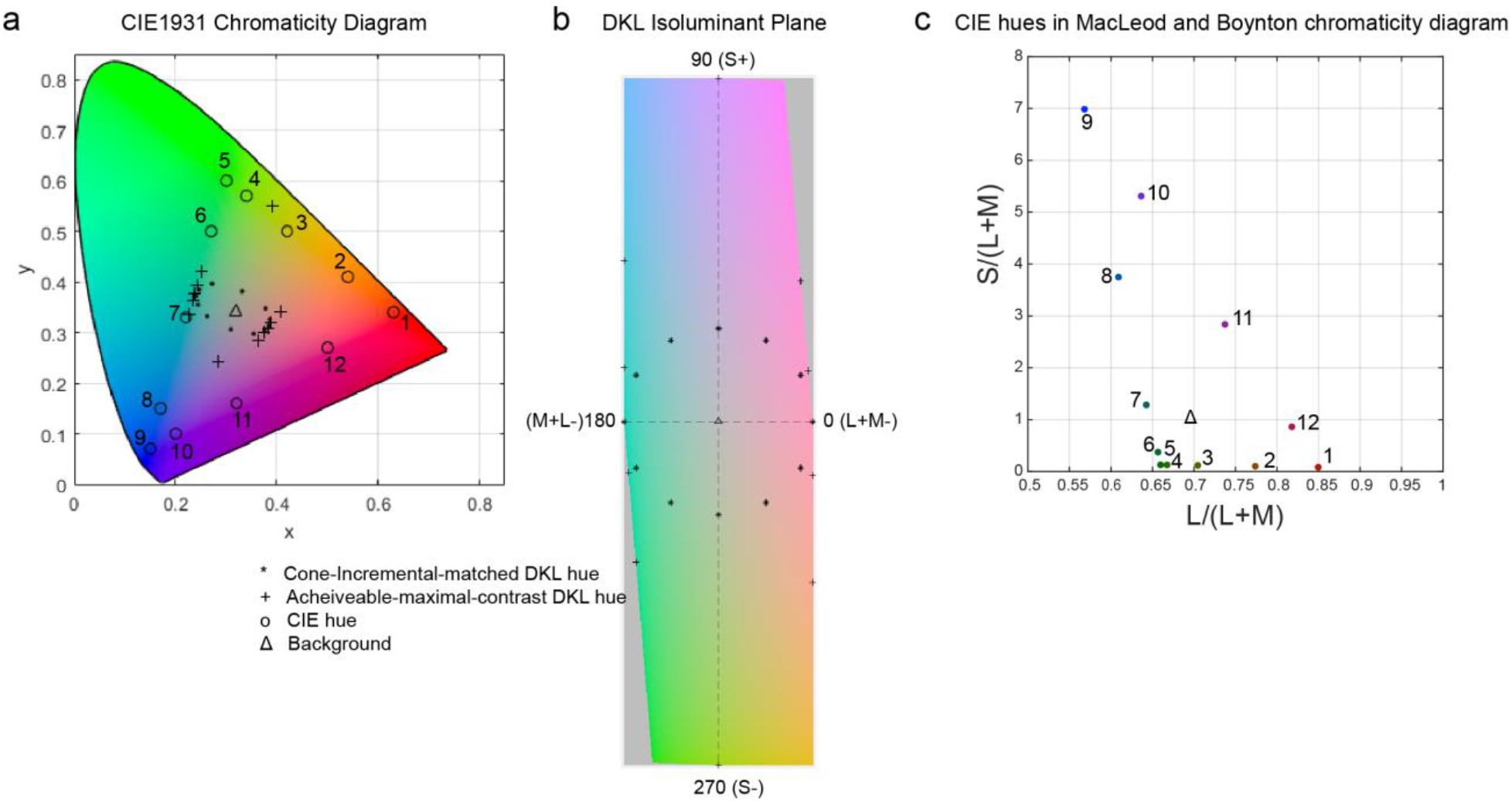
DKL and CIE hues applied in this research. **a**, Cone-increment-matched DKL hue (12 hues used in A1), achievable maximal contrast DKL hue (12 hues used for A2), and CIE hues (12 hues used in A3 and A4) are shown in CIE 1931 chromaticity diagram. Twelve CIE hues were labeled with numbers (1-12) to refer the same hue replotted in **c**. **b**, Cone-increment-matched DKL hue and achievable maximal contrast DKL hue are shown DKL isoluminant plane. **c**, Twelve CIE hues were replotted in Macleod and Boynton chromaticity diagram.

**Extended Data Fig. 9.**
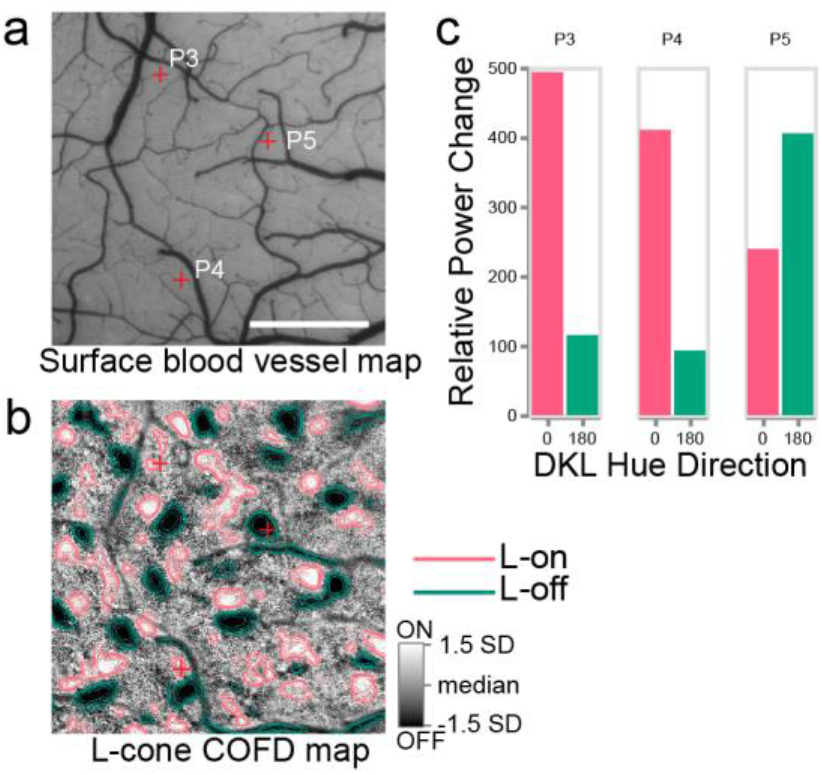
L-cone COFD map of animal A2 and electrophysiological recordings in COFDs. **a-b**, Surface blood vessel map (**a**) and L-cone COFD map (**b**) for guiding electrode penetration within COFDs. Three penetrations (red-cross) were shown in (**a-b**). Scale bar in **a**: 1 mm, applies to **a-b**. **c**, The relative change of LPF power histograms from three penetrations shows the dominant response to DKL hue at 0 degree (P3 and P4) or 180 degree (P5).

**Extended Data Table 1.** Case summary from seven animals reported in this research.

| Animal ID | Contributions to Fig. | ISI Results | 2PCI Results | 2PCI Planes | Ephys | CO |
| --- | --- | --- | --- | --- | --- | --- |
| A1 | 2 to 5, Extended Data 1, and 5 | COFD & DKL hue-phase maps, Orientation map | Cone and achromatic ON/OFF map, DKL Hue map, Cone weight distribution, Orientation map | 5 regions (10 planes) | None | Yes |
| A2 | 4, 5, Extended Data 5 and 7 | COFD & DKL hue-phase maps, Orientation map | None |  | Yes | Yes |
| A3 | 5 and Extended Data 1 | None | Cone and achromatic ON/OFF map, CIE Hue map, Cone weight distribution | 3 regions (5 planes) | None | None |
| A4 | 5 and Extended Data 1 | None | Cone and achromatic ON/OFF map, CIE Hue map, Cone weight distribution | 1 region (1 plane) | None | None |
| A5 | 6, Extended Data 3 and 5 | COFD maps, Red/Green and Blue/Yellow color maps | None |  | Yes | Yes |
| A6 | Extended Data 3 and 5 | COFD maps | None |  | None | Yes |
| A7 | Extended Data 3 and 5 | COFD maps | None |  | None | Yes |

**Extended Data Table 2.** CIE coordinates of presented hues.

| CIE Hue |  |  |  | Cone-increment-matched<br>DKL Hue |  |  |  | Achievable-maximal-contrast<br>DKL Hue |  |  |  |
| --- | --- | --- | --- | --- | --- | --- | --- | --- | --- | --- | --- |
| Hue Name | CIE Coordinates |  |  | Hue Direction<br>(degree) | CIE Coordinates |  |  | Hue Direction<br>(degree) | CIE Coordinates |  |  |
| | $x$ | $y$ | $Y_{(cd/m^2)}$ | | $x$ | $y$ | $Y_{(cd/m^2)}$ | | $x$ | $y$ | $Y_{(cd/m^2)}$ |
| Red | 0.63 | 0.34 | 10.03 | 0 | 0.36 | 0.29 | 71.54 | 0 | 0.38 | 0.31 | 85.90 |
| Orange | 0.54 | 0.41 | 10.03 | 30 | 0.36 | 0.29 | 70.99 | 30 | 0.38 | 0.30 | 85.78 |
| Yellow | 0.42 | 0.50 | 10.00 | 60 | 0.34 | 0.27 | 69.92 | 60 | 0.36 | 0.28 | 85.13 |
| Lime | 0.34 | 0.57 | 9.91 | 90 | 0.29 | 0.28 | 70.06 | 90 | 0.28 | 0.24 | 82.61 |
| Green | 0.30 | 0.60 | 9.89 | 120 | 0.24 | 0.31 | 70.07 | 120 | 0.23 | 0.34 | 84.74 |
| Green | 0.27 | 0.50 | 10.23 | 150 | 0.22 | 0.33 | 70.43 | 150 | 0.23 | 0.36 | 85.06 |
| Cyan | 0.22 | 0.33 | 10.01 | 180 | 0.22 | 0.35 | 70.39 | 180 | 0.24 | 0.38 | 85.46 |
| Blue | 0.17 | 0.15 | 9.96 | 210 | 0.23 | 0.36 | 70.85 | 210 | 0.24 | 0.39 | 85.64 |
| Blue | 0.15 | 0.07 | 10.08 | 240 | 0.25 | 0.37 | 71.51 | 240 | 0.25 | 0.42 | 86.57 |
| Purple | 0.20 | 0.10 | 9.97 | 270 | 0.31 | 0.35 | 71.83 | 270 | 0.39 | 0.55 | 89.11 |
| Magenta | 0.32 | 0.16 | 9.96 | 300 | 0.36 | 0.32 | 71.84 | 300 | 0.41 | 0.34 | 87.19 |
| Red | 0.50 | 0.27 | 9.99 | 330 | 0.37 | 0.30 | 71.49 | 330 | 0.39 | 0.32 | 86.63 |
| Background | 0.33 | 0.33 | 10.01 | Background | 0.30 | 0.31 | 71.46 | Background | 0.32 | 0.34 | 86.29 |

## Notes

### Competing Interest Statement

The authors have declared no competing interest.

## References

1. Stockman, A. & Brainard, D. H. in OSA handbook of optics (ed M. Bass) Ch. 11, 11.11–11.104 (McGraw-Hill, 2010).

2. Judd, D. B. Fundamental studies of color vision from 1860 to 1960. Proc Natl Acad Sci U S A 55, 1313–1330, doi:10.1073/pnas.55.6.1313 (1966).

3. Judd, D. B. Response functions for types of vision according to the Muller theory. J Res Natl Bur Stand (1934) 42, 1–16, doi:10.6028/jres.042.001 (1949).

4. Muller, G. E. Über die Farbenempfindungen. Zeitschrift für Psychologie und Physiologie der Sinnesorgane, Ergänzungsband 17, 1–430 (1930).

5. De Valois, R. L. & De Valois, K. K. A multi-stage color model. Vision Res 33, 1053–1065 (1993).

6. Dacey, D. M. Parallel pathways for spectral coding in primate retina. Annu Rev Neurosci 23, 743–775, doi:10.1146/annurev.neuro.23.1.743 (2000).

7. Wool, L. E., Packer, O. S., Zaidi, Q. & Dacey, D. M. Connectomic Identification and Three-Dimensional Color Tuning of S-OFF Midget Ganglion Cells in the Primate Retina. J Neurosci 39, 7893–7909, doi:10.1523/JNEUROSCI.0778-19.2019 (2019).

8. Field, G. D. et al. Functional connectivity in the retina at the resolution of photoreceptors. Nature 467, 673–677, doi:10.1038/nature09424 (2010).

9. Chatterjee, S. & Callaway, E. M. Parallel colour-opponent pathways to primary visual cortex. Nature 426, 668–671, doi:10.1038/nature02167nature02167 [pii] (2003).

10. Lennie, P., Krauskopf, J. & Sclar, G. Chromatic mechanisms in striate cortex of macaque. J Neurosci 10, 649–669 (1990).

11. Horwitz, G. D. & Hass, C. A. Nonlinear analysis of macaque V1 color tuning reveals cardinal directions for cortical color processing. Nat Neurosci 15, 913–919, doi:10.1038/nn.3105 (2012).

12. Kuriki, I., Sun, P., Ueno, K., Tanaka, K. & Cheng, K. Hue Selectivity in Human Visual Cortex Revealed by Functional Magnetic Resonance Imaging. Cereb Cortex 25, 4869–4884, doi:10.1093/cercor/bhv198 (2015).

13. Wachtler, T., Sejnowski, T. J. & Albright, T. D. Representation of color stimuli in awake macaque primary visual cortex. Neuron 37, 681–691, doi:S0896627303000357 [pii] (2003).

14. Komatsu, H., Ideura, Y., Kaji, S. & Yamane, S. Color selectivity of neurons in the inferior temporal cortex of the awake macaque monkey. J Neurosci 12, 408–424 (1992).

15. Xiao, Y., Wang, Y. & Felleman, D. J. A spatially organized representation of colour in macaque cortical area V2. Nature 421, 535–539, doi:10.1038/nature01372nature01372 [pii] (2003).

16. Tanigawa, H., Lu, H. D. & Roe, A. W. Functional organization for color and orientation in macaque V4. Nat Neurosci 13, 1542–1548, doi:10.1038/nn.2676 (2010).

17. Conway, B. R., Moeller, S. & Tsao, D. Y. Specialized color modules in macaque extrastriate cortex. Neuron 56, 560–573, doi:DOI 10.1016/j.neuron.2007.10.008 (2007).

18. Hurvich, L. M. & Jameson, D. An opponent-process theory of color vision. Psychol Rev 64, Part 1, 384–404, doi:10.1037/h0041403 (1957).

19. Hering, E. GRUNDZÜGE DER LEHRE VOM LICHTSINN. (Springer, 1920).

20. De Valois, R. L., De Valois, K. K., Switkes, E. & Mahon, L. Hue scaling of isoluminant and cone-specific lights. Vision Res 37, 885–897 (1997).

21. Mollon, J. D. & Jordan, G. in John Dalton’s Colour Vision Legacy (ed I.M.D.C.C. Dickinson) 381–392 (Taylor and Francis, 1997).

22. Krauskopf, J. in Color Vision: From Genes to Perception (eds K.R. Gegenfurtner & L.T. Sharpe) 303–317 (Cambridge University Press, 1999).

23. Livingstone, M. S. & Hubel, D. H. Anatomy and physiology of a color system in the primate visual cortex. J Neurosci 4, 309–356 (1984).

24. Ts’o, D. Y. & Gilbert, C. D. The organization of chromatic and spatial interactions in the primate striate cortex. J Neurosci 8, 1712–1727 (1988).

25. Landisman, C. E. & Ts’o, D. Y. Color processing in macaque striate cortex: electrophysiological properties. J Neurophysiol 87, 3138–3151, doi:10.1152/jn.00957.1999 (2002).

26. Lu, H. D. & Roe, A. W. Functional organization of color domains in V1 and V2 of macaque monkey revealed by optical imaging. Cereb Cortex 18, 516–533, doi:10.1093/cercor/bhm081 (2008).

27. Xiao, Y., Casti, A., Xiao, J. & Kaplan, E. Hue maps in primate striate cortex. Neuroimage 35, 771–786, doi:S1053-8119(06)01160-8[pii]10.1016/j.neuroimage.2006.11.059 (2007).

28. Landisman, C. E. & Ts’o, D. Y. Color processing in macaque striate cortex: relationships to ocular dominance, cytochrome oxidase, and orientation. J Neurophysiol 87, 3126–3137 (2002).

29. Garg, A. K., Li, P., Rashid, M. S. & Callaway, E. M. Color and orientation are jointly coded and spatially organized in primate primary visual cortex. Science 364, 1275–1279, doi:10.1126/science.aaw5868 (2019).

30. Chatterjee, S., Ohki, K. & Reid, R. C. Chromatic micromaps in primary visual cortex. Nat Commun 12, 2315, doi:10.1038/s41467-021-22488-3 (2021).

31. Ringach, D. L., Sapiro, G. & Shapley, R. A subspace reverse-correlation technique for the study of visual neurons. Vision Res 37, 2455–2464, doi:S0042-6989(96)00247-7 [pii] (1997).

32. Conway, B. R. Spatial structure of cone inputs to color cells in alert macaque primary visual cortex (V-1). J Neurosci 21, 2768–2783 (2001).

33. Conway, B. R. & Livingstone, M. S. Spatial and temporal properties of cone signals in alert macaque primary visual cortex. J Neurosci 26, 10826–10846, doi:26/42/10826 [pii]10.1523/JNEUROSCI.2091-06.2006 (2006).

34. Johnson, E. N., Hawken, M. J. & Shapley, R. The orientation selectivity of color-responsive neurons in macaque V1. J Neurosci 28, 8096–8106, doi:28/32/8096 [pii]10.1523/JNEUROSCI.1404-08.2008 (2008).

35. Horton, J. C. & Hubel, D. H. Regular patchy distribution of cytochrome oxidase staining in primary visual cortex of macaque monkey. Nature 292, 762–764 (1981).

36. Kalatsky, V. A. & Stryker, M. P. New paradigm for optical imaging: temporally encoded maps of intrinsic signal. Neuron 38, 529–545 (2003).

37. Derrington, A. M., Krauskopf, J. & Lennie, P. Chromatic mechanisms in lateral geniculate nucleus of macaque. J Physiol 357, 241–265 (1984).

38. Blasdel, G. G. & Salama, G. Voltage-sensitive dyes reveal a modular organization in monkey striate cortex. Nature 321, 579–585, doi:10.1038/321579a0 (1986).

39. Li, P. et al. A motion direction preference map in monkey V4. Neuron 78, 376–388, doi:10.1016/j.neuron.2013.02.024 (2013).

40. Obermayer, K. & Blasdel, G. G. Geometry of orientation and ocular dominance columns in monkey striate cortex. J Neurosci 13, 4114–4129 (1993).

41. Bonhoeffer, T. & Grinvald, A. Iso-orientation domains in cat visual cortex are arranged in pinwheel-like patterns. Nature 353, 429–431, doi:10.1038/353429a0 (1991).

42. Bonhoeffer, T. & Grinvald, A. The layout of iso-orientation domains in area 18 of cat visual cortex: optical imaging reveals a pinwheel-like organization. J Neurosci 13, 4157–4180 (1993).

43. Shmuel, A. & Grinvald, A. Coexistence of linear zones and pinwheels within orientation maps in cat visual cortex. Proc Natl Acad Sci U S A 97, 5568–5573, doi:97/10/5568 [pii] (2000).

44. Lu, H. D., Chen, G., Tanigawa, H. & Roe, A. W. A motion direction map in macaque V2. Neuron 68, 1002–1013, doi:10.1016/j.neuron.2010.11.020 (2010).

45. Shmuel, A. & Grinvald, A. Functional organization for direction of motion and its relationship to orientation maps in cat area 18. J Neurosci 16, 6945–6964 (1996).

46. Gegenfurtner, K. R. & Kiper, D. C. Color vision. Annu Rev Neurosci 26, 181–206, doi:10.1146/annurev.neuro.26.041002.131116041002.131116 [pii] (2003).

47. Livingstone, M. S. & Hubel, D. H. Thalamic inputs to cytochrome oxidase-rich regions in monkey visual cortex. Proc Natl Acad Sci U S A 79, 6098–6101 (1982).

48. Liu, Y. et al. Hierarchical Representation for Chromatic Processing across Macaque V1, V2, and V4. Neuron 108, 538–550 e535, doi:10.1016/j.neuron.2020.07.037 (2020).

49. Mollon, J. D. A neural basis for unique hues? Curr Biol 19, R441–442; author reply R442-443, doi:10.1016/j.cub.2009.05.008 (2009).

50. Koulakov, A. A. & Chklovskii, D. B. Orientation preference patterns in mammalian visual cortex: a wire length minimization approach. Neuron 29, 519–527, doi:10.1016/s0896-6273(01)00223-9 (2001).

51. Callaway, E. M. & Wiser, A. K. Contributions of individual layer 2-5 spiny neurons to local circuits in macaque primary visual cortex. Vis Neurosci 13, 907–922 (1996).

52. Yabuta, N. H. & Callaway, E. M. Functional streams and local connections of layer 4C neurons in primary visual cortex of the macaque monkey. J Neurosci 18, 9489–9499 (1998).

53. Lachica, E. A., Beck, P. D. & Casagrande, V. A. Parallel pathways in macaque monkey striate cortex: anatomically defined columns in layer III. Proc Natl Acad Sci U S A 89, 3566–3570 (1992).

54. Wiser, A. K. & Callaway, E. M. Ocular dominance columns and local projections of layer 6 pyramidal neurons in macaque primary visual cortex. Vis Neurosci 14, 241–251 (1997).

55. Wiser, A. K. & Callaway, E. M. Contributions of individual layer 6 pyramidal neurons to local circuitry in macaque primary visual cortex. J Neurosci 16, 2724–2739 (1996).

56. Briggs, F. & Callaway, E. M. Layer-specific input to distinct cell types in layer 6 of monkey primary visual cortex. J Neurosci 21, 3600–3608, doi:21/10/3600 [pii] (2001).

57. Leventhal, A. G., Thompson, K. G., Liu, D., Zhou, Y. & Ault, S. J. Concomitant sensitivity to orientation, direction, and color of cells in layers 2, 3, and 4 of monkey striate cortex. J Neurosci 15, 1808–1818 (1995).

58. Sadakane, O. et al. Long-Term Two-Photon Calcium Imaging of Neuronal Populations with Subcellular Resolution in Adult Non-human Primates. Cell Rep 13, 1989–1999, doi:10.1016/j.celrep.2015.10.050 (2015).

59. Jun, J. J. et al. Fully integrated silicon probes for high-density recording of neural activity. Nature 551, 232–236, doi:10.1038/nature24636 (2017).

60. Stockman, A. & Sharpe, L. T. The spectral sensitivities of the middle- and long-wavelength-sensitive cones derived from measurements in observers of known genotype. Vision Res 40, 1711–1737 (2000).

61. Brainard, D. H. in Human Color Vision 2nd Edition (eds P.K. Kaiser & R. M. Boynton) 563–579 (Optical Society of America, 1996).

62. Gatto, R. & Jammalamadaka, S. R. The generalized von Mises distribution. Statistical Methodology 4, 341–353, doi:10.1016/j.stamet.2006.11.003 (2007).

63. Swindale, N. V. Orientation tuning curves: empirical description and estimation of parameters. Biol Cybern 78, 45–56 (1998).

64. Ringach, D. L. et al. Spatial clustering of tuning in mouse primary visual cortex. Nat Commun 7, 12270, doi:10.1038/ncomms12270 (2016).

65. Ringach, D. L., Shapley, R. M. & Hawken, M. J. Orientation selectivity in macaque V1: diversity and laminar dependence. J Neurosci 22, 5639–5651, doi:2002656722/13/5639 [pii] (2002).

66. Mazurek, M., Kager, M. & Van Hooser, S. D. Robust quantification of orientation selectivity and direction selectivity. Front Neural Circuits 8, 92, doi:10.3389/fncir.2014.00092 (2014).

67. Purushothaman, G., Khaytin, I. & Casagrande, V. A. Quantification of Optical Images of Cortical Responses for Inferring Functional Maps. J. Neurophysiol. 101, 2708–2724, doi:10.1152/jn.90696.2008 (2009).

68. Park, M. & Pillow, J. W. Receptive field inference with localized priors. PLoS Comput Biol 7, e1002219, doi:10.1371/journal.pcbi.1002219 (2011).

69. Smith, S. L. & Hausser, M. Parallel processing of visual space by neighboring neurons in mouse visual cortex. Nat Neurosci 13, 1144–1149, doi:10.1038/nn.2620 (2010).

70. Niessing, J. et al. Hemodynamic signals correlate tightly with synchronized gamma oscillations. Science 309, 948–951, doi:10.1126/science.1110948 (2005).

